# Selectivity of Complex Coacervation in Multi-Protein Mixtures

**DOI:** 10.1101/2024.04.02.587643

**Authors:** So Yeon Ahn, Allie C. Obermeyer

## Abstract

Liquid-liquid phase separation of biomolecules is increasingly recognized as relevant to various cellular functions, and complex coacervation of biomacromolecules, particularly proteins, is emerging as a key mechanism for this phenomenon. Complex coacervation is also being explored as a potential protein purification method due to its potential scalability, aqueous operation, and ability to produce a highly concentrated product. However, to date most studies of complex coacervation have evaluated the phase behavior of a binary mixture of two oppositely charged macromolecules. Therefore, a comprehensive understanding of the phase behavior of complex biological mixtures has yet to be established. To address this, a panel of engineered proteins was designed to allow for quantitative analysis of the complex coacervation of individual proteins within a multi-component mixture. The behavior of individual proteins was evaluated using a defined mixture of proteins that mimics the charge profile of the *E. coli* proteome. To allow for direct quantification of proteins in each phase, spectrally separated fluorescent proteins were used to construct the protein mixture. From this quantitative analysis, we observed that the coacervation behavior of individual proteins in the mixture was consistent with each other, which was distinctive from the behavior when each protein was evaluated in a single-protein system. Subtle differences in biophysical properties between the proteins became noticeable in the mixture, which allowed us to elucidate parameters for protein complex coacervation. With this understanding, we successfully designed methods to enrich a range of proteins of interest from a mixture of proteins.

## I. Introduction

Recombinant proteins represent an important and growing class of materials in a variety of fields ranging from food, fuel, clothing, and therapeutics in a variety of clinical areas.^1–4^ Despite a growing demand for this class of biomolecules, downstream protein purification techniques have not evolved significantly from traditional methods such as liquid chromatography and precipitation.^5^ As a result, downstream protein purification is often a bottleneck in the manufacturing process and accounts for 50–80% of the overall production costs.^6^ There are a few main challenges in protein purification: the chemical similarity of the target protein and contaminating biomolecules, the necessity of mild aqueous processing conditions, and the high concentration needed for the final formulation.^7^

Simultaneously, proteins have evolved for selective enrichment within the complex cellular environment in membraneless bodies known as biomolecular condensates. Taking inspiration from the selective intracellular phase separation of proteins, one potential method to purify proteins from complex biological mixtures relies on differences in phase behavior due to varied electrostatic interactions with polyelectrolytes (PEs). This approach uses complex coacervation to selectively encapsulate the protein of interest in the coacervate phase.^8–10^ Complex coacervation, or associative liquid-liquid phase separation (LLPS) of polymers, is a phenomenon where a mixture of two oppositely charged PEs separate into dilute and dense liquid phases.^11–14^ Complex coacervation is driven by both favorable electrostatic interactions between oppositely charged PEs and bound counterion and water release.^15^ The complexation of oppositely charged PEs is entropically favorable due to the release of bound counterions and is spontaneous at modest macromolecule concentrations and moderate ionic strengths.^16^ The favorability of phase separation is directly affected by the electrostatic properties of each component such as net charge or charge identities of PEs, ionic strength of the solution, and pH.^17^ This understanding is largely derived from experimental and theoretical examination of polymer-polymer coacervates. There is more limited understanding of coacervate behavior when one of the PEs is a globular protein.^18^ But, there are boundless opportunities for the complex coacervation of proteins to enhance our understanding of coacervation and in biotechnological applications as proteins are sequence-defined polypeptides with the potential for tunable interactions with PEs.^19–21^

Employing complex coacervation as a protein purification method provides several potential advantages. First, the operation is simple with minimal to no requirements for instrumentation or labelling. Complex coacervation can be induced between oppositely charged globular proteins and polymers *in vitro*.^22,23^ Long-range non-specific electrostatic interactions provide robustness as a protein purification method. With the support of state-of-the-art studies on biomolecular condensation and accumulated knowledge of associative phase separation from polymer and colloidal science, preliminary demonstrations of the potential of complex coacervation for protein purification have been reported.^8–10,24^

Second, complex coacervation provides a soft separation method that is fully reversible. As protein function is dependent on its folding state, which can be easily disrupted, precipitation methods often result in loss of protein activity or precipitates that are difficult to dissolve.^25,26^ However, coacervates can form and dissolve reversibly as they are held together by a network of weak and transient interactions and maintain a high water content (>70%) even in the condensed phase.^8,27,28^ The physicochemical property of the dense phase can be tuned−from thermodynamically arrested solids or gels to equilibrated viscous liquids −by changing the ionic strength of the solution or chemistry of the PEs.^29,30^ Proteins that partition into the coacervate phase return to a homogenous solution successfully with no change in the folding state or enzyme activity.^31,32^ Therefore, the fact that proteins are completely soluble in both immiscible phases, rather than aggregating or precipitating, highlights the potential of complex coacervation as a promising soft protein purification method.

Lastly, highly concentrated protein can be retrieved from the dense phase, over a wide range of starting concentrations of the protein of interest. When solutions of macromolecules undergo liquid-liquid phase separation, they condense into a dense phase with macromolecule concentrations orders of magnitude higher than in the dilute phase. If the majority of the protein of interest partitions into the coacervate phase, this allows for not only selective separation of the protein of interest, but also concentration of the protein of interest. Additionally in the two-phase region of the phase diagram, changes in the initial protein fraction only affects the relative volume fractions of the dense and dilute phases.^13^ This allows a wide range of initial protein concentrations to be condensed, enhancing the protein concentration in the coacervate phase by approximately 100-fold.^25^ The dense phase can be sedimented easily by centrifugation, separated from the dilute phase and dissolved by increasing the ionic strength of the solution.

The primary goal of this study was to elucidate the governing parameters for the LLPS of globular proteins within a mixture to better understand how protein charge and sequence impact the phase behavior. Unlike complex coacervation between two linear polymers, complex coacervation with a globular protein cannot directly be predicted by charge neutrality from the isoelectric point calculated using the pK_a_s of the amino acid monomers.^33^ The distribution of charged residues, the identity of the charged amino acids, and structure-dependent solvent accessibility can affect the coacervation behavior. Herein, we focused on the dependence of complex coacervation on various charge properties of globular proteins, such as net charge and distribution of the charged residues. Frequently, LLPS is monitored by measuring the scattered light from mesoscale assemblies by an optical density measurement via a turbidity assay.^34^ Although turbidity assays detect phase separation, they do not provide any quantitative information about the composition of the dilute and coacervate phases, which are crucial to understand the phase behavior of complex mixtures and ultimately to purify proteins of interest. To address this limitation, five spectrally separated fluorescent proteins of different net charge were used to create a mixture that mimics the charge profile of the *E. coli* proteome. Distinct excitation and emission of the proteins enabled precise quantification of individual proteins in the mixture and evaluation of the phase separation behavior in a collective manner. While there have been a few attempts to selectively purify proteins through complex coacervation, these studies have primarily focused on isolating a single protein in the supernatant or from a simple mixture of two proteins.^8–10,24^ The behavior of multiple proteins, in particular the potential for coordinated behavior within a complex mixture is still not well understood. In this study, we observed independent yet coordinated behaviors of globular proteins in a mixture and establish guidelines and parameters to selectively purify a range of proteins of interest.

**Scheme 1.**
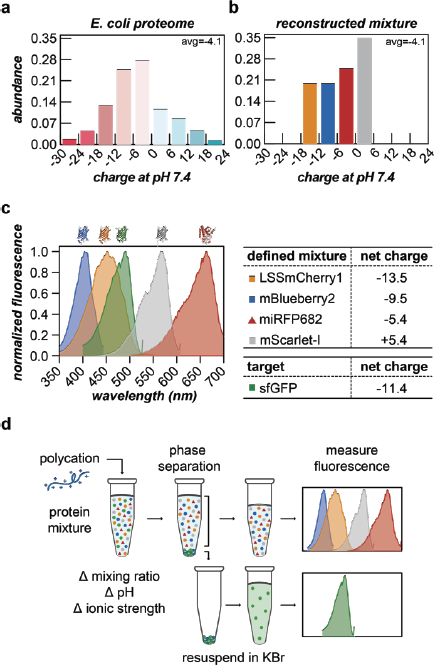
Design of defined multi-protein mixture and characterization of complex coacervation behavior. (a) Charge profile of *E. coli* proteome. (b) Charge profile of four proteins that mimic the *E. coli* proteome. Three proteins with different negative charges, Cherry-13, Blue-9 and RFP-5, were chosen to represent the negatively charged portion of the *E. coli* proteome with evenly separated charges. One positively charged protein, Scarlet+5, was chosen to represent the entirety of the positively charged portion of the proteome. The average net charge of the proteome and the reconstructed mixture were designed to be equal, with an average net charge of -4.1 (c) Absorbance spectra and net charge at pH 7.4 of fluorescent proteins used in the protein mixture. (d) Schematic of multi-protein mixture complex coacervation and characterization. Phase separation was induced by complex coacervation of the protein mixture and a polycation. The resulting two phases were physically separated through centrifugation. Proteins in each phase were quantified through selective fluorescence signals. The dense phase was redissolved with 2 M KBr (aq) before fluorescence measurement.

## II. Result & Discussion

### Fluorescent protein mixture as a model for the *E. coli* proteome

In order to quantitatively simulate the phase separation behavior of cell lysate, a complex mixture of thousands of biomolecules, a defined mixture of proteins that mimics the charge profile of the *E. coli* proteome was constructed. Protein net charge was chosen as the parameter to define the mixture, as it is one of the most dominant factors that determines the complex coacervation of proteins.^34^ Using the PaxDB protein abundance database,^35^ the charge distribution of the proteins in *E. coli* at pH 7.4 was determined (Scheme 1a). A large majority, 73%, of the proteins were negatively charged at pH 7.4, so we sought to develop an approach to purify anionic proteins using complex coacervation from a mixture that was biased towards negatively charged proteins. Four fluorescent proteins with net charge between -14 to +6, which comprise 70% of the *E. coli* proteome, were chosen to construct the mixture (Scheme 1b). To create this panel of proteins, four spectrally separated fluorescent proteins were selected: LSSmCherry1,^36^ mBlueberry2,^37^ mScarlet-I,^38^ and miRFP682 (Scheme 1c).^39^ The net charge on these proteins was then altered as needed to generate variants with distinct net charge that spanned this range of the proteome (Supplementary Table 1, Supplementary Figure 1). The mixing ratio between the four proteins was determined using the following criteria: (1) mScarlet (+5) (Scarlet+5) as the only positively charged protein represents the entirety of positively charged proteins in the proteome, accounting for approximately 30%; (2) LSSmCherry1 (-13) (Cherry-13) represents the proteins with charge below -12, approximately 20%; (3) the average net charge of the proteome and the reconstructed mixture were designed to be equal, with an average net charge of -4.1 (Scheme 1b). In addition to this mixture of four proteins, a fifth target protein of interest was added. This protein, based on superfolder GFP, had moderate anionic charge of -11 and was added to the mixture to 15 mol% of the total protein.

In order to precisely quantify the composition of the mixture using fluorescence spectroscopy, proteins that were spectrally distinct were selected. Yet when quantified using the peak excitation and emission wavelengths there was still significant overlap in the signal with > 30% interference between the samples (Scheme 1c). By careful selection of both the excitation and emission wavelengths, less than 4% of signal interference between the engineered proteins could be obtained in the proteome-mimicking mixture (Supplementary Figure 2).

### Complex coacervation of proteins in a mixture

With the protein mixture fixed, we next selected a polycation to promote complex coacervation. Initially, several polycations, including poly(allylamine hydrochloride) (PAH), quaternized poly (4-vinyl pyridine) (qP4VP), poly-L-lysine, and poly-L-arginine, were screened for the ability to selectively phase separate with proteins in the mixture. The dilute and dense phase were separated by centrifugation and proteins in each phase were quantified through fluorescence (Scheme 1d). The coacervate phase was redissolved with a high ionic strength solution (2 M KBr) for accurate spectroscopic analysis of the solubilized protein. All four polymers tested successfully prompted phase separation of the mixture of five proteins, but with differences in the partitioning of the proteins (Supplementary Figure 3). Based on these initial studies, PAH was selected for further characterization due to the high encapsulation efficiency of the anionic proteins (Supplementary Figure 4). By focusing on a single polycation, we were able to evaluate how protein properties influenced the complex coacervation behavior. Following screening, we then fixed the macromolecule concentration at 2.5 mg/mL for optimal protein encapsulation (Supplementary Figure 5).

With the total macromolecule concentration fixed, we next evaluated how the ratio of the oppositely charged macromolecules impacted complex coacervation as phase separation is optimal when the charges are balanced. To monitor the influence of the charge ratio on phase separation in a heterotypic system,^40^ the proteins and PAH were mixed at various charge ratios at a fixed total macromolecule concentration. The negative charge fraction (*f*^−^) was calculated as below, where *x* corresponds to the mass fraction of proteins, either individually or in the mixture, and *M*^−^ and *M*^+^ correspond to the charge per mass of proteins and polycation.

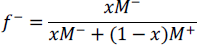

For simplicity, in the protein mixture, the weighted average charge and mass were used to calculate *M*^−^. *M*^−^ and *M*^+^ were calculated by applying the Henderson-Hasselbalch equation to the isolated amino acid side chains and the monomer of PAH at a given pH.^41^ An *f*^−^ value of 0.5 denotes charge neutrality, a state where the number of charged species are balanced in the system.

For each individual protein with PAH the maximum partitioning in the coacervate significantly deviated from predicted charge neutrality (*i.e.* charge fractions far from 0.5) (Figure 1a and Supplementary Figure 6). The charge fraction range with peak partitioning of each protein was compared by evaluating the charge fractions where the partitioning of protein in the coacervate was over 95% 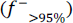 with respect to the initial input (Figure 1b). By comparing this peak partitioning metric, we see that the peak charge fraction monotonically decreased as the absolute charge of the anionic proteins decreased (*e.g.* the proteins were less negatively charged). This is somewhat counterintuitive as this indicates that more cationic polymer is required to achieve charge neutrality with less anionic proteins. However, the proteins that are less net negatively charged can undergo more induced charging upon complexation with the PAH polycation which may account for the unexpected observation.

**Figure 1.**
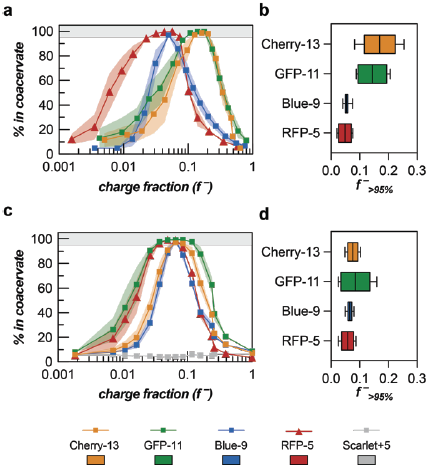
Complex coacervation of individual proteins and proteins in a mixture. (a) Partitioning of individual proteins in the coacervate phase at various charge fractions. (b) Range of charge fractions where partitioning in the dense phase is over 95% for individual proteins. (c) Coacervate phase partitioning of proteins in a mixture at various charge fractions. (d) Range of charge fraction where coacervate phase partitioning is >95% for proteins in the mixture. Total macromolecular concentrations were maintained at 2.5 mg/mL in both cases. For (a, c), data points indicate the average (a,c) and shaded regions indicate the standard error of mean (SEM). For (b, d), the box represents the 25-75 percentile with the central line indicating the median and error bars indicate the upper and lower extreme values.

### Protein complex coacervation synchronizes in a mixture

The charge faction at which maximum encapsulation was observed was different for each of the proteins when mixed independently with PAH. However, when all 5 proteins were combined with PAH the peak charge fraction was found to change, converging around *f*^−^ ∼ 0.06 (Figure 1c). This is further evident when comparing 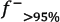 ranges. While the peak charge fraction, 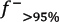, ranges minimally overlap with each other for the coacervation of individual proteins, the ranges completely overlap for the protein mixture. In the mixture, all proteins were effectively recovered (>95%) from the coacervate phase between *f*^−^ = 0.057 and *f*^−^ = 0.074 (Figure 1d). This convergent behavior of proteins in the mixture suggests that proteins not only independently interact with the polymer, but can also influence each other and alter the overall complex coacervation behavior. In order to elucidate the driving force and predict the complex coacervation behavior of a multicomponent system, it is important to understand the difference in behavior between individual proteins and the mixture and what drives components of the mixture to behave differently.

Notably, the widths of the partitioning curves also vary significantly as a function of the protein species in the mixture (Figure 1c). The width of the curve corresponds to the size of the phase separation window, an important element to consider for the selective partitioning of proteins in the coacervate. However in the protein mixture, the breadth of the phase separation window did not correlate linearly with the protein net charge, which was hypothesized to be the most influential parameter dictating electrostatic interactions with the polymer and ultimately complex coacervation behavior. Significant differences in the phase separation behavior within the mixture amplified subtle and difficult-to-quantify biophysical properties of proteins (see below), which could have been easily overlooked from comparison of individual phase behavior.

We have shown that proteins in a mixture behave in synchronous manner, but the partitioning windows do not necessarily follow a trend with respect to the protein net charge as we initially anticipated. In fact, prior work suggests that proteins with equal isoelectric point can show large differences in actual threshold pH for complex coacervation due to charge anisotropy^8^. The behavior of this protein mixture emphasizes that protein coacervation is not ruled by simple consideration of isoelectric points or net charge, but subtle differences in surface properties can lead to distinct behaviors as we observed here. For example, the peak partitioning window for GFP-11 was the broadest and encompassed that of all other proteins, followed by RFP-5, Cherry-13 and Blue-9 (Figure 1d).

### Phase behavior of the target protein is conserved across diverse mixture compositions

As cellular composition can vary from batch to batch, mixtures with varying compositions were tested in order to probe how sensitive the observed phase behavior was to fluctuations in the mixture composition. In addition, this provided corroboration of the unexpected partitioning of GFP-11 and RFP-5 within the mixture. First, mixtures with either equal molar or equal mass fraction of the five proteins were tested to evaluate if the particular mixing ratio of the proteins had a significant influence on the phase behavior of the mixture (Supplementary Figure 7). The relative partitioning of proteins was conserved independent of the precise mixing ratio tested. Analysis of the phase separation windows, via comparison of the area under the curves (AUC) analysis, showed these altered mixtures had similar relative partitioning of each protein, indicating that the distinct behaviors in the mixture are not significantly impacted by modest changes in the composition of the protein mixture (Figure 2a).

Beyond these modest composition changes, the fraction of GFP-11 was varied widely to both mimic potential purification scenarios where expression of target proteins varies in practice^42^ and to further probe the broadened phase boundary of GFP-11 in the mixture. The molar fraction of GFP-11 in the protein mixture was varied from 0 to 0.6, while maintaining a constant ratio of the remaining four proteins. AUC analysis was done with absolute molar partitioning curves (Figure 2b and Supplementary Figure 8) as well as relative partitioning curves (Figure 2c and Supplementary Figure 9). As the molar fraction of GFP-11 increased from 0 to 0.6, the AUC from the GFP-11 molar partitioning curve increased linearly, while the AUC of rest of the proteins decreased accordingly (Figure 2b). Markedly, the relative partitioning curves of individual proteins and the AUC values (Figure 2c) were preserved, regardless of the fraction of GFP-11 in the mixture. The relative protein partitioning behavior was unaffected by the mixing ratio, even when one of the five components was completely excluded. From this conserved behavior of the relative protein partitioning, we concluded that coacervation behavior of proteins was largely dependent on the property of individual proteins. Finally, we note that when initially selecting a polycation, we observed more favorable partitioning of GFP-11 and RFP-5 regardless of the polycation chemistry, further indicating that it was a property of the individual proteins.

Partitioning of individual proteins in the mixture was insensitive to extreme changes in mixing ratio and maintained the intrinsic phase boundaries, independent of the protein net charge. In order to investigate the inherent phase boundaries further, we varied the charge patterning of one of the proteins in the mixture. Charged residues on GFP-11, which were isotropically distributed across the beta-barrel, were relocated to a C-terminal peptide tag, while maintaining a similar net charge at pH 7.4. This redistribution of charged residues could potentially be used as a way to purify any protein of interest that is decoupled to the intrinsic electrostatic properties of the native protein. GFP-tag and GFP-11 individually showed distinct phase boundaries with the polycation PAH, similarly to what was reported previously by Kapelner et al^22^ (Figure 2d). However, in the proteome mimicking mixture, GFP-tag showed nearly identical phase behavior as GFP-11 (Figure 2e). This suggests the potential of charged tags to tune or enhance complex coacervation based purification of proteins with low net charge globular domains.^43,44^ Differences in the individual behavior of these GFPs suggests the importance of multivalent interactions as the isotropic distribution of charged residues is hypothesized to broaden the phase separation window due to the ability to make distinct interactions at multiple sites on the protein surface. However, in the mixture, GFP-tag, where half of the negative charge is localized to a single charge “patch” and therefore lacks additional interaction sites, behaved identically to GFP-11, indicating there were other factors at play in the mixture.

**Figure 2.**
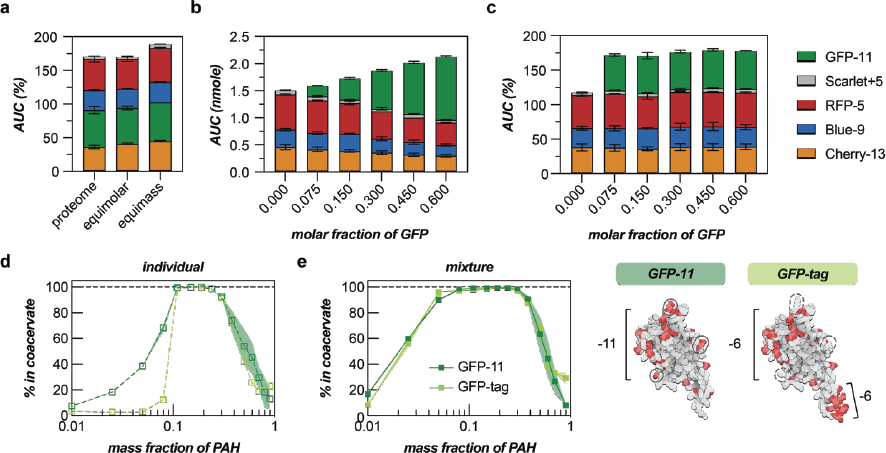
(a) AUC analysis of the relative coacervate partitioning curves of proteins in proteome mimicking mixture, equal molar mixture and equal mass mixture. (b) AUC analysis of the absolute partitioning curves with varied fraction of GFP-11 in the mixture. (c) AUC analysis of the relative partitioning curves with respect to initial input of proteins. The fraction of Cherry-13, Blue-9, RFP-5 and Scarlet+5 was constant in each of the mixtures. (d) Partitioning of GFP variants in the coacervate phase at various mass fractions. (e) Coacervate phase partitioning of GFP variants in the proteome mimicking mixture. (right) Residues in red indicate anionic residues, Asp and Glu. Residues on GFP-11 circled by filled lines were substituted with neutral or cationic residues on GFP-tag, circled by dashed lines. Total macromolecular concentrations were maintained at 2.5 mg/mL. Shaded regions and error bars indicate the SEM.

### Scaffold-Client model

Within a mixture of polyelectrolytes, such as the one studied here, molecules can either drive coacervate formation or simply partition into a coacervate phase formed by other PEs. This phenomenon is commonly observed in biomolecular condensates that form in the complex intracellular environment in the presence of thousands of biomolecules. To simplify some of the complexity of multi-component phase behavior, a scaffold and client model has been adopted in this field, where scaffolds are essential for condensate formation and clients favorably partition into the condensed phase. Similar effects have been observed in synthetic systems, as shown by Blocher et al. where incorporation of a client protein shifted the pattern of complex coacervation between two scaffold PEs.^33^ In the protein mixture studied here we observe that proteins can participate as clients at conditions where individually they do not phase separate (Supplementary Figure 10a). A shift in the partitioning between the individual protein and the mixture was especially noticeable for Blue-9, where the peak PAH mass fraction migrated from 0.37 to 0.19 (Supplementary Figure 6 a and b). This can potentially be attributed to proteins acting as clients at specific conditions when the coacervation is led by other proteins. Interestingly, GFP-11 showed significantly broadened partitioning in the mixture, encompassing that of all other anionic proteins, in addition to migration of the partitioning window (Supplementary Figure 10b). In other words, GFP-11 appeared to more readily participate in complex coacervation as a client as compared to the other anionic proteins.

**Table 1.** Protein parameters relevant to complex coacervation. Protein net charge was predicted using monomeric pK_a_ values and PROPKA3 based on the predicted 3D structure of the protein.^45^ The 3D structure of each protein was predicted using ColabFold.^46^ The Grand Average of Hydropathy (GRAVY) was calculated as the sum of the hydropathy values for all the amino acids in the protein divided by the total number of residues.^47^ Positive hydropathy index values are assigned for more hydrophobic residues. Solvent accessible area was calculated by EDTsurf and excluded the common disordered HisTag on the N-terminus of each protein.^48^ *<u>Underlined and italicized values</u>* indicate properties that are differentiated from the rest of the proteins.

|  | Calculation | Cherry-13 | GFP-11 | Blue-9 | RFP-5 |
| --- | --- | --- | --- | --- | --- |
| **net charge at pH 7.4** | linear | -13.57 | -11.41 | -9.57 | -5.40 |
|  | PROPKA | -12.13 | -10.93 | -8.43 | -5.66 |
| **isoelectric point** | linear | 5.15 | 5.64 | 5.58 | 6.25 |
|  | PROPKA | 5.82 | 5.67 | 6.19 | 6.30 |
| **total hydrophobic residues**<br>(Ala, Val, Ile, Leu, Met, Phe, Tyr, Trp) |  | 82 | 84 | 87 | <u>140</u> |
| **GRAVY score** |  | -0.78 | -0.60 | -0.68 | <u>-0.04</u> |
| **total charged residues** |  | 70 | <u>62</u> | 68 | 68 |
| Asp + Glu |  | 42 | 37 | 39 | 37 |
| Arg + Lys |  | 28 | 25 | 29 | 31 |
| negative charge density ($\frac{Asp+Glu}{total\ residues}$) | | 0.17 | 0.15 | 0.16 | 0.11 |
| ratio of negative-to-positive residues |  | 1.5 | 1.5 | 1.3 | 1.2 |
| **solvent accessible area ($\text{\AA}^2$)** |  | 12181.97 | 11557.04 | 12425.24 | <u>16348.34</u> |

With this observation, we asked: Why does GFP most favorably partition into complex coacervates? Among the proteins with the same beta-barrel structure, (Cherry-13, GFP-11, Blue-9, and Scarlet+5), GFP-11 displayed the broadest partitioning window despite having intermediate net charge (Figure 1c). These three proteins were largely similar, with nearly identical length, molecular weight, structure, and surface area, but with different expected net charge. Given the unexpected behavior of GFP-11, further examination of the sequences and possible biophysical differences was warranted. First, we noticed that there were fewer charged residues, both negative and positive, on GFP-11 than Cherry-13 and Blue-9 (Table 1). Indeed, GFP-11 and Cherry-13 have nearly the same negative-to-positive ratio as defined by Obermeyer et al^34^ (Table 1) despite the net charge difference. GFP-11 showed the most favorable partitioning behavior and had the lowest number of charged residues among the anionic proteins, potentially indicating that the relative distribution of charged residues and the net charge of patchy regions impacts the phase behavior as has been seen in computational studies.^49^

We next evaluated how the specific folded structure impacted the net charge to further probe possible differences between the proteins that could be responsible for deviations from expectations. PROPKA3 was used to predict the pK_a_ values of individual residues based on the folded structure of proteins predicted by ColabFold.^45^ The pH dependent net charge of Cherry-13 and Blue-9 was overestimated when the 3D protein structure was not taken into account, while that of GFP-11 and RFP-5 were less dependent on the folded state of the protein (Supplementary Figure 11a, b, c and d). This pattern was clearer for the isoelectric points, with the folded GFP-11 showing the lowest value of 5.64 (Supplementary Figure 11e). These deviations were consistent whether evaluated at the whole protein level or at the individual residue level and all indicated that GFP-11 was more likely to be negatively charged than Cherry-13 or Blue-9 (Supplementary Figure 11f, g, h, I, j, and k). This suggests that GFP-11, which has a higher ionization potential and lower pKa, can more favorably induce additional negative charge when in contact with the polycation, further increasing the chance for partitioning in the coacervate.

Then it comes down to one question: why is the net charge of Cherry-13 and Blue-9 overestimated while GFP-11 is not? Although they share a beta-barrel structure, Cherry-13, GFP-11 and Blue-9 have different locations and relative distances between charged residues, which can result in distinct charged states, ionization potentials, and potential for charge regulation upon PE complexation. To probe this, we analyzed approximate distances between like and oppositely charged residues to potentially explain the distinct protonation state of GFP-11. We found that like charges were closely distributed for Cherry-13 and Blue-9 compared to GFP-11 and RFP-5 (Supplementary Figure 12a). Close proximity of similar charges results in a high-energy repulsive situation, altering the pK_a_ of individual residues. On the other hand, the distances between opposite charges were closer for GFP-11 such that salt bridge formation could occur, with distances <4 Å (Supplementary Figure 12b). The heterogenous distribution of charges on structurally rigid proteins affected the protonation state, leading certain proteins to be more prone to charge regulation. Protein residue analysis also revealed that RFP-5 significantly deviated from expectations based on net charge.

However, this protein was the only one in the mixture that did not adopt a beta-barrel structure common to many fluorescent proteins,^50^ and this completely different structure, size, and pattern of surface potential also deviated most from the other proteins. Notably, while RFP-5 is negatively charged, RFP-5 also has more hydrophobic residues and a higher GRAVY score compared to the other anionic proteins in the mixture (Table 1, Supplementary Figure 14). Protein condensation has been shown to be impacted by both electrostatic and hydrophobic interactions.^51–53^ We propose that the interplay of electrostatics, which drive the phase transition, and hydrophobicity, which can serve to stabilize the condensed phase, broadens the partitioning window of this protein. In addition, the larger surface area of RFP-5 could be advantageous for the formation of multivalent interactions with the PE.

These deviations from expectations were seemingly coupled to differences in the physical properties of the coacervates as characterized by microscopy as well (Supplementary Figure 13). Coacervates of GFP-11 and PAH as well as RFP-5 and PAH, both of which had the broadest partitioning efficiency, appeared to be liquid-like. On the other hand, coacervates between PAH and Cherry-13 or Blue-9, which had narrower partitioning windows, displayed fiber-or solid-like properties at the same mass fraction where GFP-11 and RFP-5 formed liquid-like coacervates. Coacervates formed from the protein mixture and PAH showed liquid-like properties at the same mass condition, indicating that the GFP and RFP interactions likely dominated and resulted in the formation of the coacervate. The correlation between the wide phase separation window and liquid-like property of coacervate droplets can be explained from the ability of proteins to maintain charge neutrality by charge regulation, facilitating network formation via weak, multivalent interactions.^54,55^

### Distinct behaviors of individual proteins in a mixture to enable selective coacervation

To further evaluate protein phase behavior as a function of protein charged states, we investigated how proteins interacted with the oppositely charged PE as a function of the solution pH. Complex coacervation between the proteome mimicking mixture and PAH was initiated at pH 6.4 to 11.4 to evaluate how the phase boundaries of individual proteins in the mixture responded to changes in pH (Figure 3a, b, c, d and Supplementary Figure 15). The two main factors predicted to dictate the phase boundaries were the charge state of the protein and of PAH, both of which have ionizable residues with varying pKa values. As the pH increases, proteins in the mixture become more negatively charged, but PAH, with a monomer pKa of 8.8^56^ becomes less positively charged. This results in decreased interaction strength between the proteins and polycation at elevated pH and results in a non-monotonic pattern with an initial increase in the AUC as the pH increases from 6.4 to ∼9.4 followed by a dramatic decrease as the pH is further increased (Figure 3e). The area under the curve for each protein was analyzed to quantitatively evaluate the partitioning behaviors as a function of pH. Near neutral pH (6.4-8.4), we observed an unexpected relationship between the protein net charge and the AUC, with the AUC decreasing in the following order GFP-11 > RFP-5 > Cherry-13 > Blue-9. As the pH was further increased, the AUC for RFP-5 decreased more quickly than the other more highly charged protein variants, such that at pH 10.4 the AUC analysis nearly followed the expected trend. While the absolute partitioning did not correlate with expected protein net charge, the peak pH values for partitioning of each protein did. The pH values at the peak AUC (pH_peak_) for each protein were determined through a spline fit to the data (Table 2) and were linearly correlated with the net charge of proteins (Figure 3f), indicating that the overall net charge of individual proteins contributes to pH dependent phase behavior with PAH.

Next, we investigated the pH response of each protein during coacervate dissolution. In contrast to the coacervate formation analysis, where the spontaneity of phase separation is evaluated, dissolution of the coacervates allows evaluation of the interaction strength with the polycation and is essential for protein recovery following coacervation. As described above, pH conditions were varied initially when the coacervates were formed by mixing the protein mixture with PAH at the pre-determined pH value (Figure 3g). In contrast, pH-driven dissolution was evaluated by exposing a coacervate formed at pH 7.4, that had been separated from the supernatant, to various pH solutions (Figure 3h). The mass fraction where all four anionic proteins partition nearly 100% in the coacervate (mass fraction of PAH 0.19) was chosen to compare how formation and dissolution of the coacervate depended on the solution pH. When pH was varied initially during the coacervate formation, the least negatively charged proteins, Blue-9 and RFP-5, showed the greatest pH sensitivity and nearly identical behavior in response to increasing pH, followed by similar behavior of the more negatively charged proteins, Cherry-13 and GFP-11 (Figure 3h). When centrifuged coacervates formed at pH 7.4 were resuspended at various pH conditions (Figure 3i), dissolution behavior for all four proteins was distinct from that of formation (Supplementary Figure 16). For dissolution, the least anionic protein, RFP-5 was more resistant to increasing pH than Blue-9, perhaps due to increased hydrophobic interactions that are not pH sensitive. Additionally, the most anionic proteins, Cherry-13 and GFP-11, were not released from the coacervate phase even at pH values that did not support coacervate formation.

**Figure 3.**
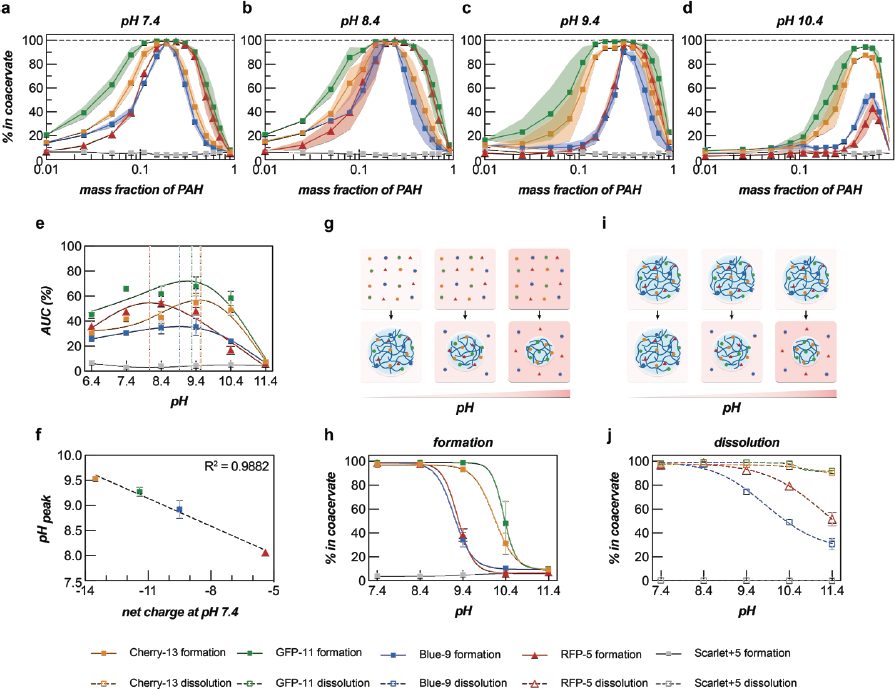
Complex coacervation of protein mixtures at various pH conditions. Partitioning of proteins in the coacervate phase with respect to the initial input quantity at (a) pH 7.4 (b) pH 8.4 (c) pH 9.4 and (d) pH 10.4. (e) AUC analysis of partitioning curves at various pH. Dotted lines indicate the pH_peak_ from the spline fit (solid lines). (f) Linear correlation between protein net charge and pH_peak_ values from the AUC analysis. (g) Schematic of coacervate formation with pH variation. (h) Partitioning of proteins in the coacervate phase as the function of pH variation during coacervation formation. A mass fraction of 0.19 PAH was chosen for coacervate formation. (i) Schematic of coacervate dissolution at varying pH conditions after coacervate formation at a fixed pH. (j) Partitioning of proteins in the coacervate phase during dissolution at various pH conditions. The coacervate was first assembled at pH 7.4, at a PAH mass fraction of 0.19 before being exposed to various pH conditions. Total macromolecular concentrations were maintained at 2.5 mg/mL. Shaded regions and error bars indicate the SEM.

In parallel, we investigated how the complex coacervation of each protein in the mixture was affected by the ionic strength of the solution. Increases in the ionic strength of the solution results in a smaller phase separation window, as electrostatic interactions are screened and the effect of entropic gain from counterion release is reduced. The ionic strength of the coacervation solution was varied by adding KBr to the solution at concentrations ranging from 0 to 800 mM KBr (Figure 4a, b, c, d and Supplementary Figure 17). From 0 mM to 50 mM KBr, the overall phase boundaries for all four anionic proteins broadened with increased AUCs (Figure 4e). This overall increase in protein partitioning in the coacervate in the presence of some exogenous counterions can be attributed to the formation of a network structure that recruits more proteins and has been observed in other complex coacervates.^12,54^ The breadth of the phase separation window, as monitored by the AUCs, of individual proteins monotonically decreased between 50 and 800 mM KBr (Figure 4e). As expected, the AUC of RFP-5 decreased more than the others with increases in the solution ionic strength. The half maximal inhibitive concentration (IC50) was estimated from a sigmoidal model fit to evaluate the response of individual proteins to increases in ionic strength. IC50 values followed the expected net charge trends, even though the absolute AUC values of proteins were inconsistent with the net charge (Table 2 and Figure 4f). This suggests that while individual proteins are sensitive to the ionic strength during coacervate formation, once the coacervate forms, proteins interact with PAH independently following their own charge. This shows that the binding of individual proteins in the coacervate phase is dictated by independent interactions with PAH, which is related to the overall protein charge.

Similar to pH, the impact of solution ionic strength on both coacervate formation and dissolution was investigated. The mass fraction where the anionic proteins maximally partitioned in the coacervate was chosen for comparison of coacervate formation and dissolution (mass fraction of PAH 0.19). Differences were observed for the salt concentrations required for dissolution and the critical salt concentration for coacervate formation (Figure 4g, h, I, j and Supplementary Figure 18). However, these differences were relatively minor compared to those observed for pH changes. This can be attributed to differences in how altering the pH and ionic strength impact coacervation. Changes in pH alter the protonation state of the protein and polymer, likely impacting the enthalpy of complexation. Conversely, as the solution ionic strength is varied there is less entropy gained from bound counterion release, thus shifting the equilibrium.

**Table 2.** pH_peak_ and IC50 values of anionic proteins in the mixture. pH_peak_ and IC50 values were determined through a spline fit and sigmoidal fit respectively to the AUC data. Estimated values represent resistance to loss in protein-PE interaction strength due to pH and ionic strength increase.

|  | pH<sub>peak</sub> | IC<sub>50</sub> (mM) |
| --- | --- | --- |
| **Cherry-13** | 9.5 | 658.5 |
| **GFP-11** | 9.3 | 636.3 |
| **Blue-9** | 8.9 | 567.1 |
| **RFP-5** | 8.1 | 499.9 |

**Figure 4.**
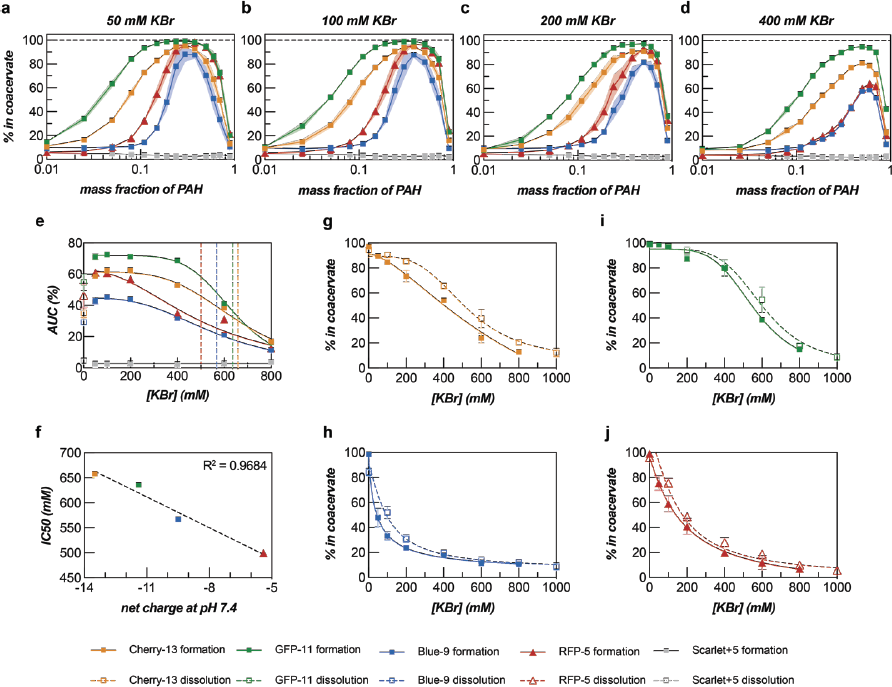
Complex coacervation of protein mixtures at various ionic strength conditions. Partitioning of proteins in the coacervate phase at KBr concentrations of (a) 50 mM (b) 100 mM (c) 200 mM and (d) 400 mM. (e) AUC analysis of partitioning curves at various ionic strengths. Data sets were fitted with a sigmoidal model between 50 and 800 mM KBr. Dotted lines indicate the IC50 for each protein from the sigmoidal model fit. (f) Linear correlation between protein net charge and IC50 values of AUC analysis. Partitioning of (g) Cherry-13 (h) GFP-11 (i) Blue-9 and (j) RFP-5 in the coacervate phase during formation and dissolution of the coacervate at various ionic strengths. A mass fraction of 0.19 PAH was chosen for coacervate formation. The coacervate was assembled at 0 mM KBr concentration before dissolution assay. Total macromolecular concentrations were maintained at 2.5 mg/mL. Shaded regions and error bars indicate the SEM.

### Protein purification via complex coacervation

With an improved understanding of how the overall net charge and charge heterogeneity govern the phase behavior of proteins in a mixture, we finally sought to separate and concentrate proteins from the mixture via complex coacervation. Using the distinct behavior of the proteins in response to the ionic strength of the solution, individual proteins were selectively separated from the mixture. Using the broad phase boundary and resistance to high ionic strength, GFP-11 was separated to 81% molar purity, which corresponds to 5.4-fold increase from the original mixture (Figure 5). This was accomplished by a few simple steps: (1) adding a polycation to the protein mixture at a specific ratio that favors phase separation of only highly anionic proteins (mass fraction of 0.19), (2) centrifuging and isolating the coacervate phase, (3) partial dissolution of the coacervate phase with modest salt concentrations (40 mM NaCl) to release contaminating proteins, and finally (4) addition of KBr to resuspend the purified GFP. This purity and recovery were achieved using the polycation qP4VP based on our initial screening of polycations and was achieved with minimal optimization after this observation. (Supplementary Figure 3). Furthermore, 65% of the initial input of the protein was retrieved while achieving the high purity separation. Purification of GFP-11 with PAH resulted in 60% molar purity with 45% of the target protein recovered, without optimization of the purification process. Likewise, Blue-9 and RFP-5 were purified to 69% and 60% on a molar basis, while 48% and 80% of the initial input were recovered, respectively. As expected, Scarlet+5 did not phase separate with the polycation individually or in the mixture, even in the case of near complete partitioning of all anionic proteins (Figure 1b). This led to superb purity and recovery of 99% and 96%, respectively, by simply recovering the supernatant following coacervation of the anionic proteins.

In comparison to standard methods for separation by protein precipitation, such as ammonium sulfate precipitation, complex coacervation showed high selectivity and softness. Following a similar separation and quantification procedure, ammonium sulfate was shown to successfully precipitate the proteins in the mixture. However, the purity of proteins either in the supernatant or the precipitate was not improved compared to the original mixture due to the similar solubilities of the proteins in the mixture (Supplementary Figure 19).^57^ Unlike complex coacervation where fine tuning of the partitioning is possible through variations in ionic strength and pH before and after coacervation, similar options for systematic manipulation were not feasible besides varying the salt concentration. In addition, there were significant protein losses during the ammonium sulfate purification process. The total recovery of proteins from both the precipitate and the supernatant was incomplete, with only 56% of the initial input recovered from both phases, which indicates compromised protein folding or solubility following precipitation. In contrast, recovery of protein from both phases was quantitative for complex coacervation with PAH, emphasizing the softness of this alternate phase separation method.

**Figure 5.**
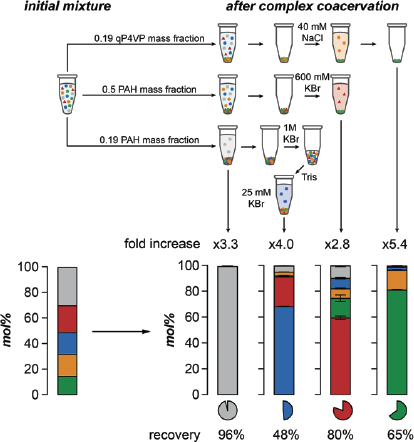
The molar purity, purity increase, and recovery of individual proteins after selective complex coacervation and separation. The molar purity of Scarlet+5, Blue-9, RFP-5 and GFP-11 are 99%, 68%, 60% and 81%. Separations were performed on the 100 µL scale with a macromolecule concentration of 2.5 mg/mL. The maximum purity of mScarlet+5 was achieved from the dilute phase at 0.19 PAH mass fraction. The maximum purity of Blue-9 was achieved by redissolving the coacervate formed at 0.19 PAH mass fraction in 2.5 µl of 1 M KBr (aq), then adding 97.5 µl of 10 mM Tris-HCl buffer to dilute the KBr (aq) and selectively re-induce coacervation. The maximum purity of RFP-5 was achieved by redissolving the coacervate formed at 0.5 mass fraction in 600 mM KBr (aq). The maximum purity of GFP-11 was accomplished with a coacervate formed at 0.19 qP4VP mass fraction and redissolved in 30 mM NaCl (aq). Shaded regions and error bars indicate the SEM.

## IV. Conclusion

In this study, the complex coacervation behavior of proteins with varying surface charge was evaluated by directly quantifying the proteins in the coacervate. In addition to characterizing the behavior of individual proteins, the use of spectrally separated fluorescent proteins also enabled the quantitation of a protein mixture. The phase behavior of individual proteins showed significant differences when in the mixture. The charge fraction where maximum partitioning was observed converged in the mixture, but the phase separation window was distinct for each protein and somewhat independent of the net charge of the proteins. As has been seen previously,^22,34,58^ the complex coacervation of globular proteins did not peak at the expected charge neutral conditions and required significant excess of the polycation. Some proteins appeared to participate as client molecules, exhibiting broadened two-phase regions encompassing that of other proteins. By monitoring the behavior of a protein mixture, subtle differences in electrostatic properties between the proteins, which was difficult to quantify *a priori*, became more noticeable. In a complex coacervation system, transient but cumulative interactions are amplified to a measurable transition, which cannot be fully understood by single isolated parameter such as net charge or hydrophobicity.

Detailed protein surface properties were monitored through coacervate formation and dissolution. The pKa values of the ionizable residues predicted based on the 3D structure were markedly lower for GFP-11, which showed the broadest phase boundary and favorable partitioning as a client. Low pKa values not only indicate the protein is more negatively charged than anticipated, but also mean that the protein can go through more extensive charge regulation upon complexation with the PE. Stronger charge regulation can tune the charge imbalance in the coacervate enabling phase separation in expanded conditions, ultimately broadening the phase boundaries. This suggests that balancing the charge in the coacervate through charge regulation, plays a key role in how proteins form complex coacervates.

While proteins showed synchronized partitioning behavior with distinct boundaries determined by charge regulation, the response to the solution ionic strength and pH remained consistent with the net charge of proteins. From monitoring the partitioning of proteins at varying ionic strength and pH during coacervate formation and dissolution, we were able to probe both the spontaneity of the protein partitioning, and the binding between the protein and the PEs. In contrast to the size of the partitioning window, the binding, as monitored by dissolution, was more consistent with traditional parameters such as net charge.

By utilizing the elucidated differences in the phase boundaries, we were able to effectively separate the target GFP-11 from the proteome mimicking mixture and achieved 81% purity, while recovering 65% of the original GFP-11 in the mixture. We look forward to further refine this method to enrich a single protein of interest from the complex mixture of proteins in the *E. coli* lysate. Moreover, we expect this evaluation of the behavior of individual proteins in a defined mixture is expected to provide guidance for other applications that require selective complex coacervation of proteins such as protein delivery and *in vivo* biomolecular condensation.

## Associated Content

The Supporting Information is available free of charge at https://xxx.xxx/xxx.

The PDF contains experimental details as well as supplementary data including: protein sequence, SDS-PAGE, LC-MS characterization of proteins, analysis of fluorescence, polycation selection, protein quantification in the supernatant and the coacervate phase, coacervate partioning optimization, coacervate phase volume measurement, partitioning analysis of various mixture compositions, pH and ionic strength conditions, PROPKA3 analysis, opposite- and like-charge distance analysis, microscopic images of phase separation, hydropathy analysis, and analysis of ammonium sulfate precipiation.

Further information and requests for resources and reagents should be directed to and will be fulfilled by the lead contact, Allie Obermeyer. Plasmids for the production of the engineered fluorescent proteins used here have been deposited with Addgene (#219415-#219424). Custom MATLAB codes for like- and opposite-charge distribution analysis are publicly available on GitHub (https://github.com/Obermeyer-Group).

## Author Information

### Author Contributions

Conceptulization, S.Y.A. and A.C.O.; Investigation, S.Y.A.; Data analysis and writing – original draft and review and editing, S.Y.A. and A.C.O.

### Notes

The authors declare the following competing financial interst(s): A.C.O. is a co-founder of Werewool, a company that is engaged in the development of performance textiles that incorporate engineered proteins. S.Y.A. declares no competing interests.

## Acknowledgments

This work was supported by the National Institutes of Health under award number NIH-NIGMS-R35GM138378.

