## Supplementary Information for "Selectivity of Complex Coacervation in Multi-Protein Mixtures"

### Table of Contents

#### Methods

Cloning 2
Protein expression 2
Protein purification 2
Protein preparation 2
Liquid chromatography mass spectrometry 2
Complex coacervation induction and separation 2
pH variation during coacervate formation and dissolution 2
Ionic strength variation during coacervate formation and dissolution 2
Coacervate phase volume measurement 3
Fluorescence microscopy 3
Individual protein quantification (recovery assays) 3
Area under curve analysis 3
Calculation of protein net charge 3
Analysis of E. coli proteome and charge abundance 3
Protein structure prediction 3
PROPKA 3

#### Supplementary Figures

Supplementary Table 1. Summary of the protein panel 4-5
Supplementary Figure 1. Analysis of purified proteins 6
Supplementary Figure 2. Fluorescence of the five proteins selected for the mixture 7
Supplementary Figure 3. Partitioning analysis of proteins in the mixture when complexed with various polycations 8
Supplementary Figure 4. Protein partitioning analysis and mass balance 9
Supplementary Figure 5. Optimization of protein partitioning condition 10
Supplementary Figure 6. Visualization of complex coacervation 11
Supplementary Figure 7. Partitioning analysis of proteome-mimicking, equimolar, equimass mixtures 12
Supplementary Figure 8. Absolute partitioning in the proteome-mimicking mixture at various GFP-11 molar fractions 13
Supplementary Figure 9. Relative partitioning in the proteome-mimicking mixture at various GFP-11 molar fractions 14
Supplementary Figure 10. Partitioning comparison of individual proteins and proteins in the mixture 15
Supplementary Figure 11. Protein net charge and $pK_a$ values of residues predicted from linear sequence and PROPKA3 16
Supplementary Figure 12. Distance analysis of negatively charged residues 17
Supplementary Figure 13. Microscopic images of the coacervates of individual proteins and protein mixture 18
Supplementary Figure 14. Hydropathy values of residues anionic proteins and 2D mapping of protein surface hydropathy 19
Supplementary Figure 15. Partitioning analysis of proteins in the mixture at various pH 20
Supplementary Figure 16. Mixture behavior during formation and dissolution of the coacervates at various pH 21
Supplementary Figure 17. Partitioning analysis of proteins in the mixture at various ionic strength 22
Supplementary Figure 18. Mixture behavior during formation and dissolution of the coacervates at various ionic strength 23
Supplementary Figure 19. Absolute recovery of proteins during ammonium precipitation 24

**Cloning.** The DNA sequences of GFP-11, with charge isotropically distributed or localized to a peptide tag, in a protein expression vector (pETDuet) were previously constructed.[1] DNA sequences of Cherry-13, Blue-9, miRFP682 (RFP-5), and Scarlet+5 cloned in pET28a (+) were purchased from TWIST Bioscience. Mutations replacing neutral (Ile, Leu, Ala, Val, Thr, Met, Asn, Gln) and basic (Arg, Lys) residues with acidic (Glu, Asp) residues on the surface of LSSmCherry1 (-9.8), sfGFP (-5.6), and mBlueberry2 (-3.8) were made to increase the overall net negative charge on the proteins to create Cherry-13, GFP-11, and Blue-9, respectively. Neutral and acidic residues on the surface of mScarlet-I were replaced with basic residues to reverse the net charge of the protein from negative to positive creating Scarlet+5. Mutation sites for GFP-11 were chosen from the set of mutations previously reported to generate a supercharged GFP with an expected charge of (-30).[2] Similarly, mutation sites for Cherry-13, Blue-9, and Scarlet+5 were chosen from highly solvent-exposed sites from crystal structures and evenly distributed across the protein surface to minimize the introduction of charge patches and disruption of protein folding[3–5].

**Protein expression.** All proteins were expressed in NiCo21(DE3) *E. coli* cells in 1 L cultures of LB media. LB media was supplemented with 100 µg of ampicillin for GFP mutants and 100 µg of kanamycin for the other fluorescent proteins. Cultures were incubated at 37 ºC, with shaking at 175 rpm. Cultures were grown to $OD_{600}$ of 0.9 and were subsequently induced by addition of 1 mL of 1 M isopropyl β-D-1-thiogalactopyranoside (IPTG). The cultures were incubated with shaking at 25 ºC for 16 h after induction.

**Protein purification.** Cells from 1 L cultures were harvested by centrifugation at 3488 g for 15 min, and cell pellets were resuspended in lysis buffer (50 mM $NaH_2PO_4$, 300 mM NaCl, pH 8.0; 15 mL buffer per 1 L of culture) and frozen at -80 ºC overnight. The thawed cells were lysed by sonication with a microtip (1/8 in, 2 s on, 4 s off, total 10 min sonication time with 40% amplitude) and cell debris was removed by centrifugation at 11,627 g for 45 min. The proteins were purified using Ni-NTA affinity chromatography as described by the manufacturer with the following procedure: (i) 7.5 mL of resin was used per 1 L of culture, (ii) the resin was equilibrated with 50 mL of lysis buffer, (iii) 15 mL of the clarified lysate was loaded to the column, (iv) protein-bound resin was washed once with 50 mL lysis buffer and twice with 50 mL wash buffer (50 mM $NaH_2PO_4$, 300 mM NaCl, 50 mM imidazole, pH 8.0), (v) protein was eluted from the resin with 50 mL of elution buffer (50 mM $NaH_2PO_4$, 300 mM NaCl, 250 mM imidazole, pH 8.0). Fractions were collected from steps (iii) through (v) and were analyzed by SDS-PAGE using a 4 –12% gradient gel.

**Protein preparation.** The pure fractions containing the protein of interest were dialyzed against 4 L of 10 mM Tris buffer (pH 7.4) using a regenerated cellulose dialysis membrane with a 3.5 kDa molecular weight cut-off (MWCO). At least seven buffer changes were performed after a minimum 3 h interval to ensure complete buffer exchange. Buffer exchanged proteins were filtered with a polyethersulfone (PES) syringe filter with a pore size of 0.22 µm to remove aggregates. Filtered proteins were concentrated by ultrafiltration using 10 kDa MWCO Amicon Ultra centrifugal filter units to a final protein concentration of 3 mg/mL. The concentration of each protein was determined by measuring the absorbance and using the reported extinction coefficient at $\lambda_{maxima}$.[3,4,6–9]

**Liquid chromatography mass spectrometry.** Sample solutions were made by diluting the protein stock solution to 0.1 mg/mL in Milli-Q purified water. Reversed-phase chromatography was performed using an Agilent 1260 HPLC system equipped with a 1 × 75 mm InifinityLab Poroshell 300Extend-C18 column (671750-902, Agilent Technologies). Solvent A and B were composed of 0.1% formic acid in Milli-Q water and acetonitrile, respectively. The flow rate was 0.4 mL/min and the column temperature was set at 25 °C. A sample volume of 5 µL was injected onto the column. The gradient elution profile was from 3 to 90% B for 4 minutes followed by washing with 97% B for 1 min. The column was equilibrated with 3% B for 10 min prior to analysis. Full mass data were acquired with a range of 100−3000 m/z and an acquisition rate of 1 spectrum/s on an Agilent 6230 LC/TOF system. The instrument was operated in positive mode for intact protein analysis. Mass spectrum was deconvoluted using Agilent MassHunter BioConfirm software with the built in Maximum Entropy algorithm.

**Complex coacervation induction and separation.** A panel of proteins−Cherry-13, Blue-9, RFP-5 and Scarlet+5 −was mixed with a molar ratio of 0.2, 0.2, 0.25, and 0.35, respectively, to mimic the charge distribution of the *E. coli* proteome. GFP variants were added to this panel at 15% of the total protein mixture, unless mentioned otherwise, as a target protein to be purified. To a small volume of Tris-HCl buffer, a 3 mg/mL solution of poly (allylamine hydrochloride) (PAH) solution (average $M_w$ 17,500, Sigma Aldrich) was added to a desired mass fraction. Subsequently, a 3 mg/mL solution of an individual protein or the protein mixture with the given ratio was mixed with the diluted polymer solution. Samples were mixed in a 96-well round bottom plate and incubated for 30 min at room temperature. The solutions were centrifuged at 3500 g for 30 min to separate the dense and dilute phases. 80 µL of the dilute phase was retrieved for measurement and the remaining supernatant (approximately 20 µL) was discarded. Then, 100 µL of a 2 M KBr solution was added to the dense phase and gently pipetted to redisperse the coacervate. After resuspension, 80 µL of this solution was removed for analysis. The separation and analysis described here were used for all experiments described below unless mentioned otherwise.

**pH variation during coacervate formation and dissolution.** For initial pH variation, 10 mM Tris-HCl buffer, protein solutions, and polymer solution were titrated to the desired pH separately prior to mixing. For pH dependent dissolution of the coacervates, the dense phase isolated by centrifugation was resuspended with 100 µL of 10 mM Tris-HCl buffer with pH varying from 6.4 to 11.4 by gentle pipetting and bath sonication (5 s on, 5 s off, repeated 3 times). The partially resuspended dense phase was incubated for 30 min at room temperature, followed by separation and analysis.

**Ionic strength variation during coacervate formation and dissolution.** For initial ionic strength variation, the KBr concentration in 10 mM Tris buffer, protein solution, and polymer solution was adjusted prior to mixing. For ionic strength dependent dissolution of the coacervates, the dense phase isolated by centrifugation was resuspended with 100 µL of a KBr solution, with KBr concentration ranging from 100 mM to 1000 mM, by gentle pipetting and bath sonication (5 s on, 5 s off, repeated 3 times). The partially resuspended dense phase was incubated for 30 min at room temperature, followed by separation and analysis.

**Coacervate phase volume measurement.** Polymer and protein solutions were mixed in PCV cell counting tubes (TP87005, MIDSCI) with a total volume of 500 µL. 30 min after mixing, the dense phase was collected in the capillary by centrifugation at 3500 g for 10 min. Dense phase that adhered to the tube wall was resuspended by sonication in a water bath (5 s on, 5 s off, repeated 6 times) followed by a second centrifugation step at 3500 g for 10 min. The PCV cells were imaged using a Perfection V850 Pro scanner (Epson) and volumes of the dense phases were estimated through two-dimensional area analysis using ImageJ.[10]

**Fluorescence microscopy.** Microscopic images were taken using a Nikon Ti Eclipse fluorescence microscope. Samples were prepared at 2.5 mg/mL total macromolecular concentration with varied mass fractions in a 384 well glass bottom plate (Cellvis P384-1.5H-N). The total volume of each sample was 50 µL. All images were taken using a 20× Plan-Apochromat NA 0.75, long working distance. Blue-9, GFP-11, Cherry-13, Scarlet+5 and RFP-5 were imaged using 395/430(40 bandwidth), 470(40)/525(50), 440/590(40), 592(8)/625(15) and 637(30)/680(42) nm excitation and emission filters respectively.

**Individual protein quantification.** Each protein was quantified using a plate reader (M200 Pro, Tecan) at a specific $\lambda_{excitation}/\lambda_{emission}$ to ensure selective quantification of the individual protein in the mixture. The following excitation and emission wavelengths were used for each protein: 656 nm/692 nm for RFP-5, 580 nm/610 nm for Scarlet+5, 470 nm/520 nm for GFP variants, 410 nm/640 nm for Cherry-13 and 392 nm/460 nm for Blue-9. At each of these selected wavelengths the fluorescence signal from the protein of interest was at least 95% of the total signal. Calibration curves were made with individual proteins with and without 2 M KBr for analysis of the supernatant and the coacervate, respectively. Fluorescence signals measured from the dilute and dense phases at specific wavelengths were used to estimate the quantity of the individual proteins in each phase.

**Area under curve analysis.** Area under the partitioning curves were computed using Area Under the Curve analysis in GraphPad Prism. The area of trapezoids defined by linear lines connecting the discrete XY values of the partitioning-mass fraction plots was summed.

**Calculation of protein net charge.** The net charge for each protein was calculated as a function of pH using a MATLAB (MathWorks Inc.) script[11] based on the Henderson-Hasselbach equation[12] and amino acid information taken from Organic Chemistry by T. W. Graham Solomons and Craig B. Fryhle.[13]

**Analysis of *E. coli* proteome and charge abundance.** Using the PaxDB 5.0 absolute protein abundance database[14] and UniProt[15] protein sequence database, the distribution of expected protein net charge as a function of cellular abundance in *E. coli* was determined. Abundance data of *E. coli* strain K-12 substrain MG1655 (Uniprot Taxonomy ID 511145) was used for analysis.

**Protein structure prediction.** Three-dimensional (3D) structures of proteins were predicted using ColabFold v1.5.2, which enables the accelerated homology search of MMseqs2 with AlphaFold2.[16,17] Predicted structures of Cherry-13, GFP-11, Blue-9 and Scarlet+5 were aligned with the 3D structure sfGFP (PDB ID: 2B3P) on Pymol v2.5.2 to confirm consistent β-barrel structure.

**PROPKA3.** Isoelectric points of individual residues on proteins were calculated with PROPKA3[18,19] using the 3D structures predicted from ColabFold.

| | net charge at pH 7.4 | sequence | modifications | peak $\lambda_{ex}/\lambda_{em}$ (optimized $\lambda_{ex}/\lambda_{em}$) (nm) | ref |
|---|---|---|---|---|---|
| **LSSmCherry1 (Cherry-13)** | -13.6 | MGHHHHHHGG VSKGEEDNMT<br>IIKEFMRFKV HMEGSVNGHE<br>FEIEGEGEGR PYEGTQTAKL<br>KVTKGGPLPF AWDILSPQFM<br>YGSKAYVKHP ADIPDYLKLS<br>FPEGFNWERV MNFEDGGVVT<br>VTQDSSLQDG EFIYKVELRG<br>TNFPSDGPVM QCRTMGLEAS<br>TERMYPEDGA LKGESKERLK<br>LKDGGHYDAE VKTTYKAKEP<br>VQLPGAYNVD IKLDILSHNE<br>DYTIVEQYER SEGRHSTGGM<br>DELYK | K137E, K199E | 450/610 (410/640) | 7 |
| **mBlueberry2 (Blue-9)** | -9.6 | MGHHHHHHGG VSKGEENNVA<br>IIKEFMRFKV HMEGSVNGHE<br>FEIEGEGEGR PYEGTQTAKL<br>KVTKGGPLPF AWDILSPQFL<br>FGSKVYIKHP ADIPDYFKLS<br>FPEGFKWERV MNFEDGGVVT<br>VTQDSSLQDG VFIYKVELRG<br>TNFPSDGPVM QKKTMGWEAF<br>SERMYPEDGA LKSEIKTRLK<br>LKDGGHYDAE VKTTYKAKEP<br>VQLPGAYNVN IKLDIVSHNE<br>DYTIVEQYER AEGEHSTGGM<br>DELYK | K137E, K199E, R234E | 402/467 (392/460) | 8 |
| **miRFP682 (RFP-5)** | -5.4 | MGHHHHHHGG AEGSVARQPD<br>LLTCDDEPIH IPGAIQPHGL<br>LLALAADMTI VAGSDNLPEL<br>TGLAIGALIG RSAADVFDSE<br>THNRLTIALA EPGAAVGAPI<br>TVGFTMRKDA GFIGSWHRHD<br>QLIFLELEPP QRDVAEPQAF<br>FRRTNSAIRR LQAAETLESA<br>CAAAAQEVRK ITGFDRVMIY<br>RFASDFSGVV IAEDRCAEVE<br>SKLGLHYPAS AVPAQARRLY<br>TINPVRIIPD INYRPVPVTP<br>DLNPVTGRPI DLSFAILRSV<br>SPCHLEFMRN IGMHGTMSIS<br>ILRGERLWGL IVCHHRTPYY<br>VDLDGRQACK RVAERVLATQ<br>IGVMEE | - | 663/682 (656/692) | 8 |
| **miRFP682 (RFP-5)** | -5.4 | MGHHHHHHGG AEGSVARQPD<br>LLTCDDEPIH IPGAIQPHGL<br>LLALAADMTI VAGSDNLPEL<br>TGLAIGALIG RSAADVFDSE<br>THNRLTIALA EPGAAVGAPI<br>TVGFTMRKDA GFIGSWHRHD<br>QLIFLELEPP QRDVAEPQAF<br>FRRTNSAIRR LQAAETLESA<br>CAAAAQEVRK ITGFDRVMIY<br>RFASDFSGVV IAEDRCAEVE<br>SKLGLHYPAS AVPAQARRLY<br>TINPVRIIPD INYRPVPVTP<br>DLNPVTGRPI DLSFAILRSV<br>SPCHLEFMRN IGMHGTMSIS<br>ILRGERLWGL IVCHHRTPYY | - | 663/682 (656/692) | 9 |

| | | | | | |
|---|---|---|---|---|---|
| | | VDLDGRQACK RVAERVLATQ IGVMEE | | | |
| **sfGFP (GFP-11)** | -11.4 | MGHHHHHHGG ASKGEELFTG VVPILVELDG DVNGHKFSVR GEGEGDATEG KLTLKFICTT GKLPVPWPTL VTTLTYGVQC FSRYPDHMKQ HDFFKSAMPE GYVQERTISF KDDGTYKTRA EVKFEGDTLV NRIELKGIDF KEDGNILGHK LEYNFNSHNV YITADKQENG IKANFKIRHN VEDGSVQLAD HYQQNTPIGD GPVLLPDNHY LSTQSALSKD PNEDRDHMVL LEFVTAAGIT HGMDELYK | N49E, K168E, K224D | 485/510 (470/520) | 2,3,20 |
| **GFP-tag** | -12.4 | MGHHHHHHGG ASKGEELFTG VVPILVELDG DVNGHKFSVR GEGEGDATNG KLTLKFICTT GKLPVPWPTL VTTLTYGVQC FSRYPDHMKQ HDFFKSAMPE GYVQERTISF KDDGTYKTRA EVKFEGDTLV NRIELKGIDF KEDGNILGHK LEYNFNSHNV YITADKQKNG IKANFKIRHN VEDGSVQLAH YQQNTPIGDG PVLLPDNHYL STQSALSKDP NEKRDHMVLL EFVTAAGITH GMDELYKDEE EDD | C-terminal DEEEDD tag | 485/510 (470/520) | 20 |

**Table 1**. Summary of the protein panel. miRFP682[9] was used unmodified. LSSmCherry1,[7] mBlueberry2,[8] mScarlet-I,[4] and sfGFP[3] were mutated to have the desired net charge while ensuring minimal impact on the protein structure using PyMol (v. 2. 5. 2.) and ColabFold[16,17]. Peak and optimized excitation and emission wavelengths correspond to spectra peak and conditions used for selective quantification of individual proteins.

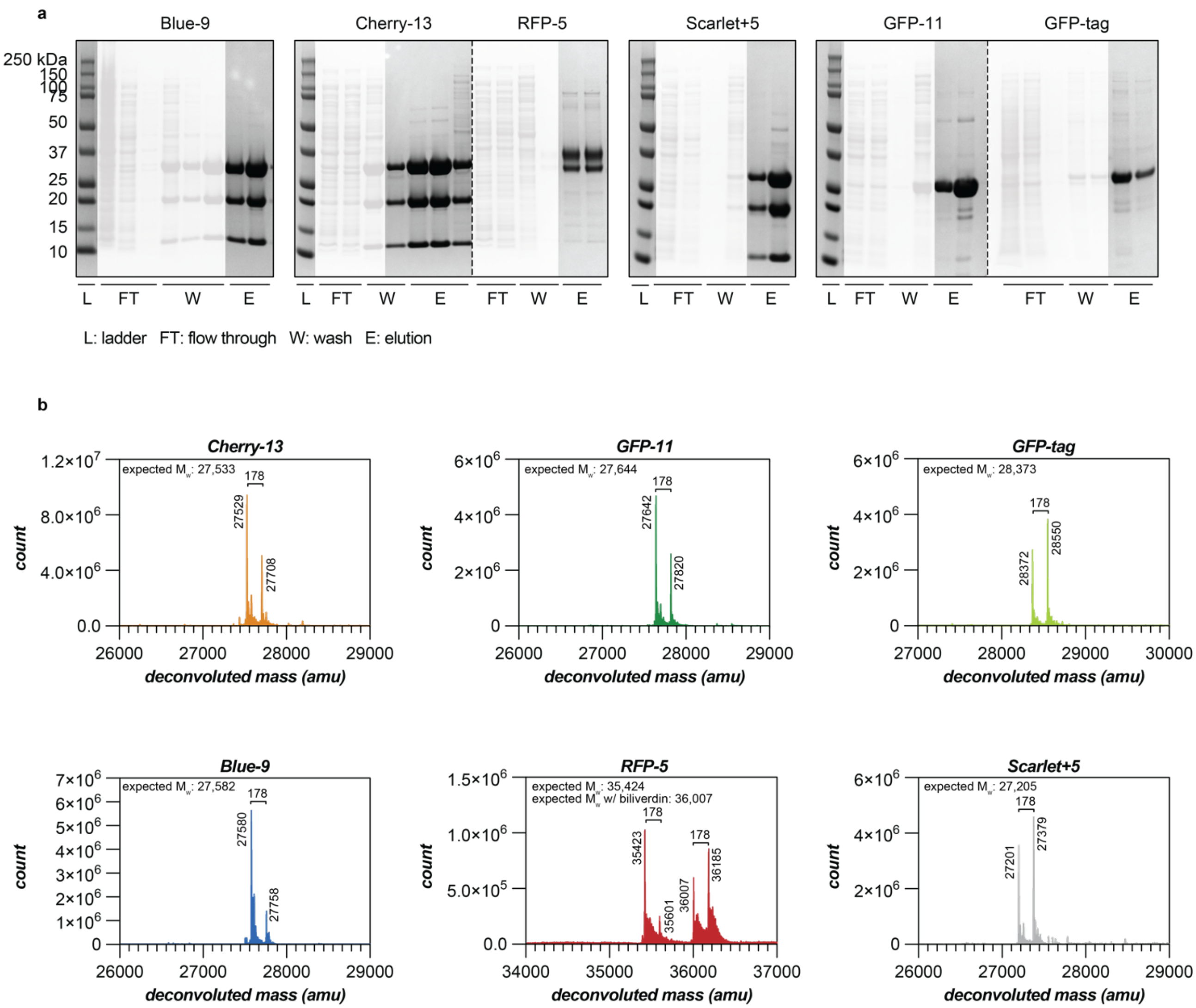


**Supplementary Figure 1**. Analysis of purified proteins. (a) SDS-PAGE analysis of the purified proteins used herein demonstrates purity after immobilized metal affinity chromatography. Impure fractions collected during purification were shaded and only the non-shaded pure fractions were used for subsequent experiments. Proteins with an imine in the chromophore, Blue-9, Cherry-13, and Scarlet+5, showed two lower molecular weight bands due to imine hydrolysis during analysis. Two bands are seen in RFP-5 lanes as RFP-5 does not fully denature by SDS[21]. (b) LC-MS analysis of the purified proteins demonstrates that all expressed proteins have the correct expected molecular weight. Expected molecular weights were determined assuming methionine truncation. The 178 Da shift from the expected molecular weight observed in all samples, corresponds to a-N-6-Phosphogluconoylation of a His-tag in *E. coli*[22].

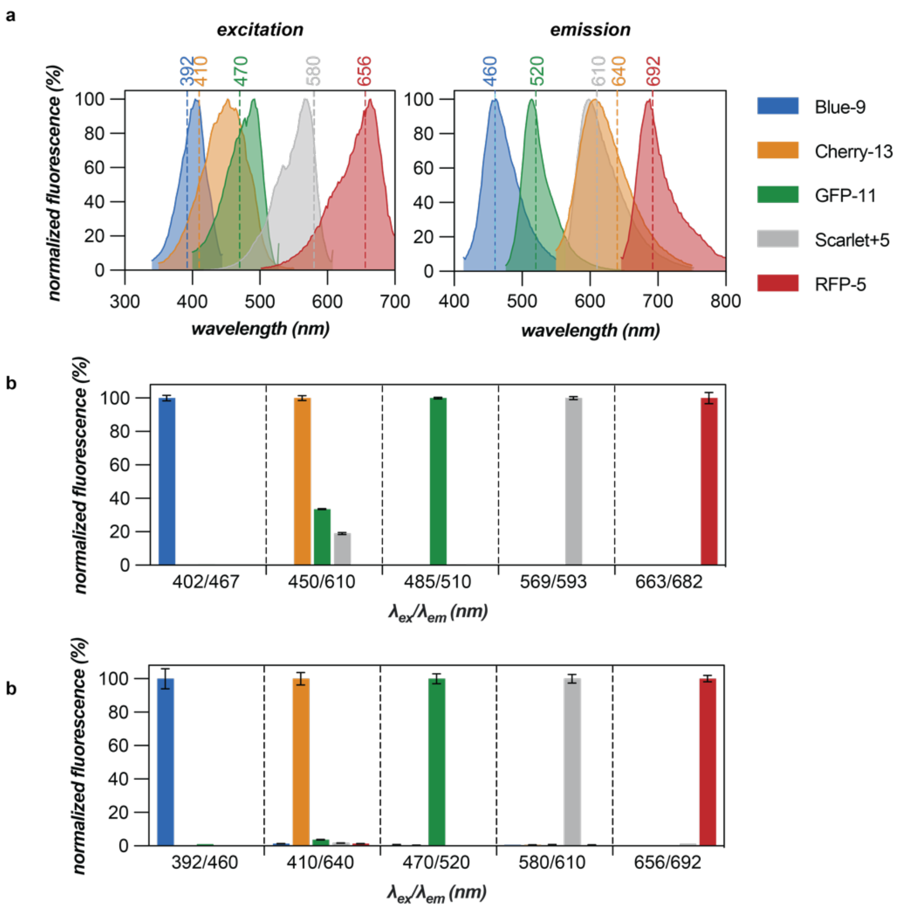


**Supplementary Figure 2**. Fluorescence of the five proteins selected for the mixture. (a) Normalized excitation and emission scans of each individual protein. The peak excitation and emission wavelengths reported in FPbase were used when measuring the excitation and emission scans, respectively[3,4,7–9]. Dotted lines indicate the excitation and emission wavelengths optimized for minimum signal interference among the proteins in this mixture. (b) The normalized fluorescence of all of the five proteins when measured at the reported peak excitation and emission wavelengths for each of the proteins. (c) The normalized fluorescence of all of the five proteins when measured at the optimized excitation and emission wavelengths for each of the proteins. The maximum interference in this case was <4%.

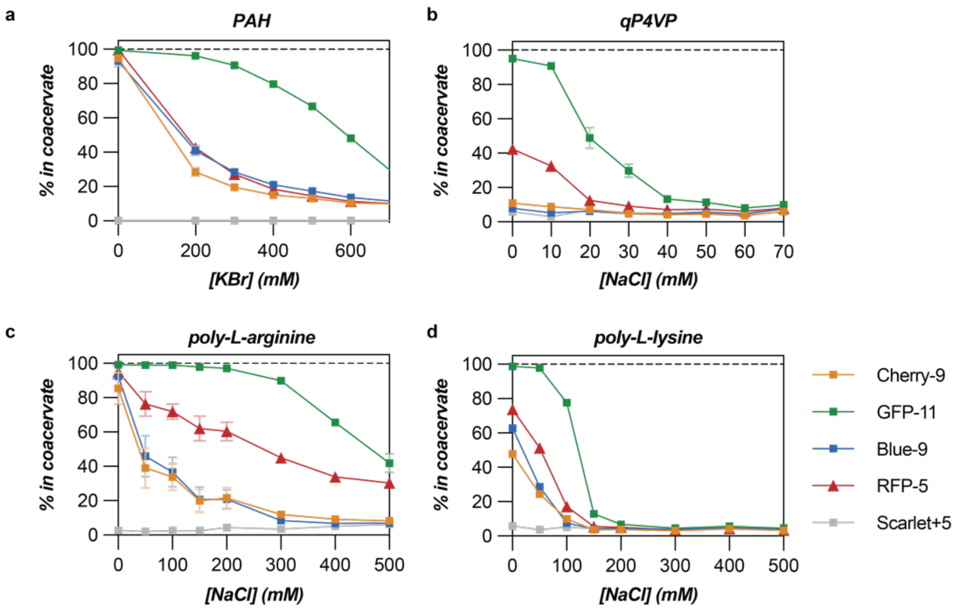


**Supplementary Figure 3**. Partitioning of proteins in the coacervate when complexed with various polycations. Complex coacervation of the protein mixture with (a) PAH, (b) qP4VP, (c) poly-L-arginine, (d) poly-L-lysine. Coacervation of a 2 mg/mL solution of the protein mixture was induced with 0.5 mg/mL polycation in 10 mM tris buffer with varying concentrations of salt (KBr (a) or NaCl (b-d)). Wildtype LSSmCherry[7] (net charge of -9.6 at pH 7.4) was used for the protein mixture instead of modified Cherry-13.

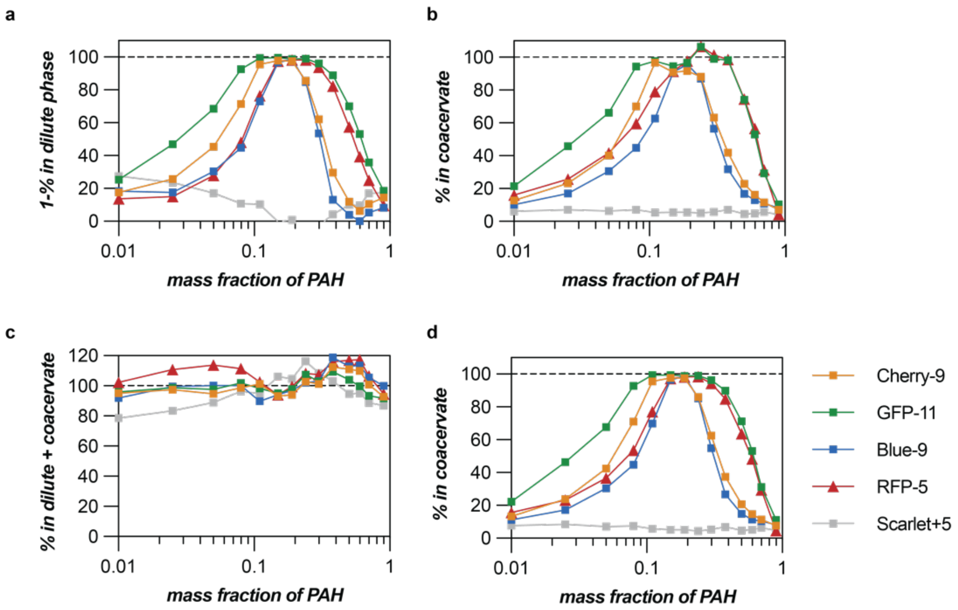


**Supplementary Figure 4**. Protein partitioning analysis. Fraction of proteins in the coacervate phase (a) estimated by subtracting the quantity in the dilute phase with respect to the initial input and (b) directly quantified from the redissolved coacervate. (c) The sum of proteins in both phases with respect to the initial input. (d) Adjusted partitioning curve with respect to the total proteins quantified for both phases. The partitioning in the coacervate was estimated by the fraction of proteins quantified in the coacervate phase from the total protein quantified in both phases. The partitioning of proteins in (a) and (b) are in reasonable agreement with the adjusted partitioning in (d), and the sum of proteins in both phases are in good agreement with the initial input of individual proteins. Adjusted partitioning curve was used for all situations for the estimation of protein partitioning across a broad range of protein and polymer concentrations. Complex coacervation was induced at total macromolecule concentration of 2.5 mg/mL, pH 7.4 with PAH.

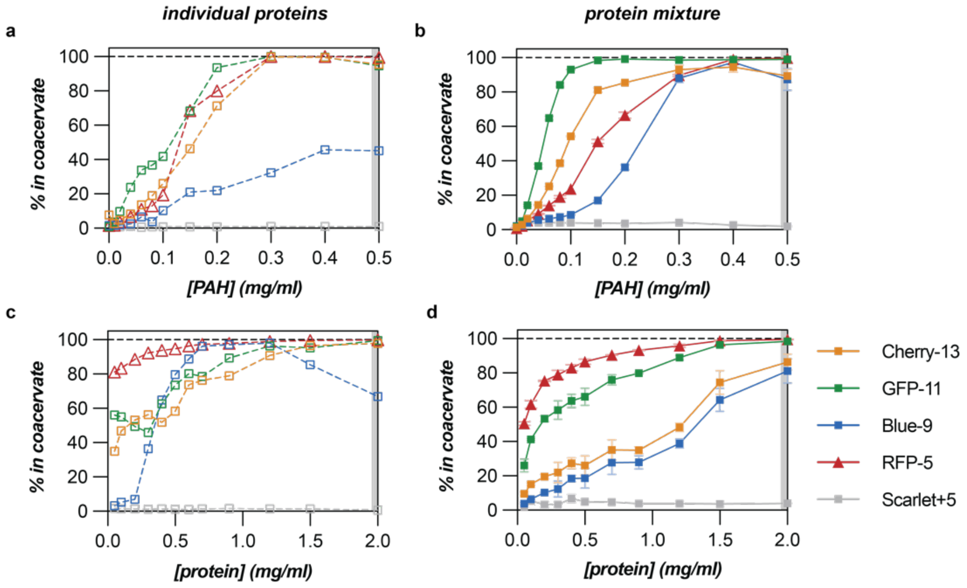


**Supplementary Figure 5**. Optimization of protein partitioning condition. PAH concentration variation with (a) 2 mg/mL individual proteins or (b) a 2 mg/mL protein mixture. (c) Individual protein concentration variation and (f) protein mixture concentration variation at a fixed 0.5 mg/mL PAH concentration. Shaded bars indicate the optimized condition.

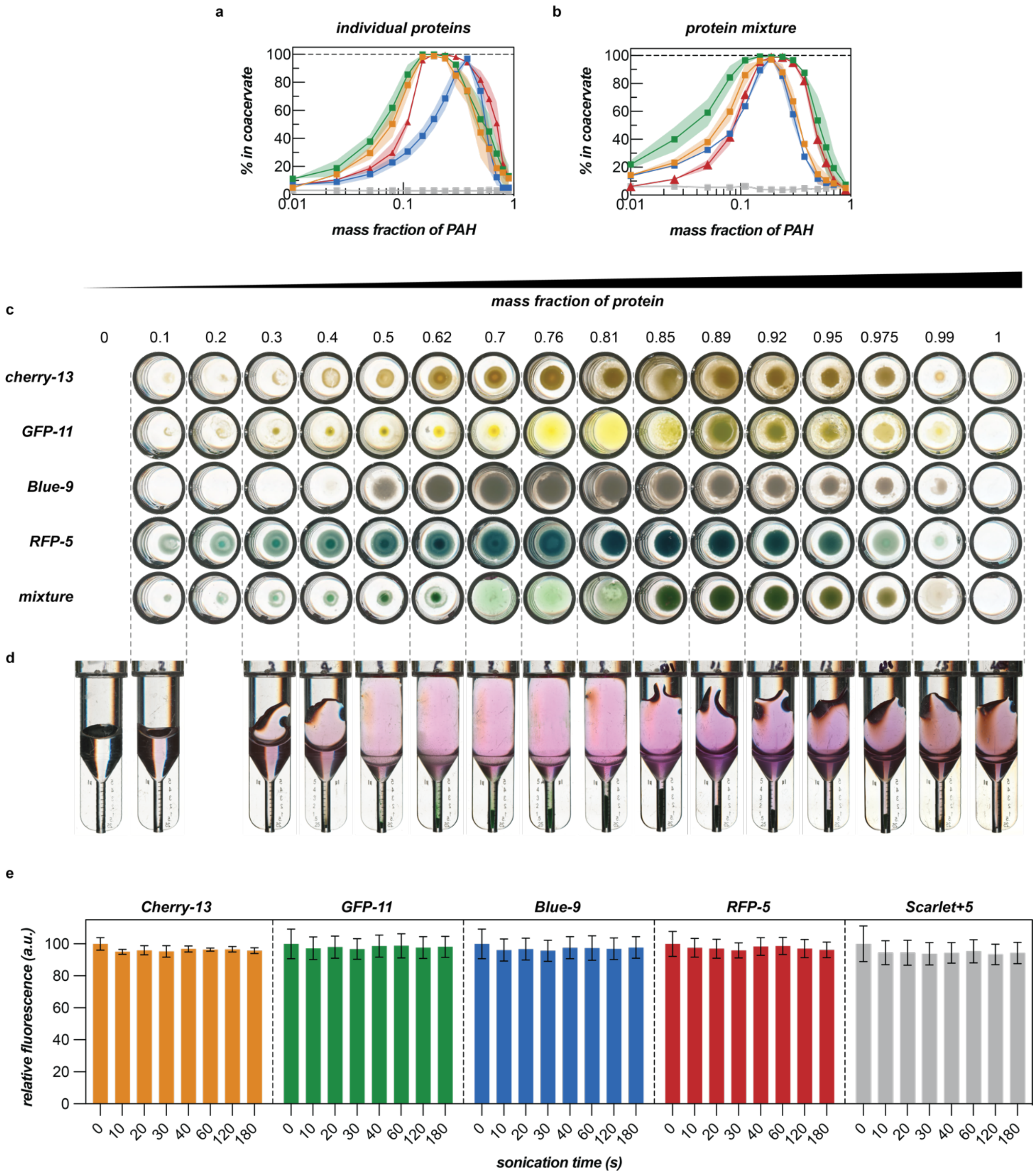


**Supplementary Figure 6**. Visualization of complex coacervation. Partitioning curve of (a) individual proteins and (b) protein mixture following complex coacervation with PAH across a range of polymer mass fractions. (c) Images of the coacervate phase in a well plate after centrifugation and decantation of the dilute phase. (d) Separated coacervate phase and dilute phase of complex coacervation of protein mixture visualized in PCV cell counting tubes. The coacervate phase was collected in the bottom capillary by centrifuging the phase separated mixture. The centrifuged samples were sonicated in a water bath to minimize adhesion of the coacervates on the wall of the tubes and then centrifuged a second time to consolidate the coacervate phase into a single large phase. (e) Relative fluorescence of individual proteins over the course of bath sonication. No fluorescence signal alteration was observed with up to 180 s of sonication. Complex coacervation was induced at total macromolecule concentration of 2.5 mg/mL, pH 7.4 with PAH.

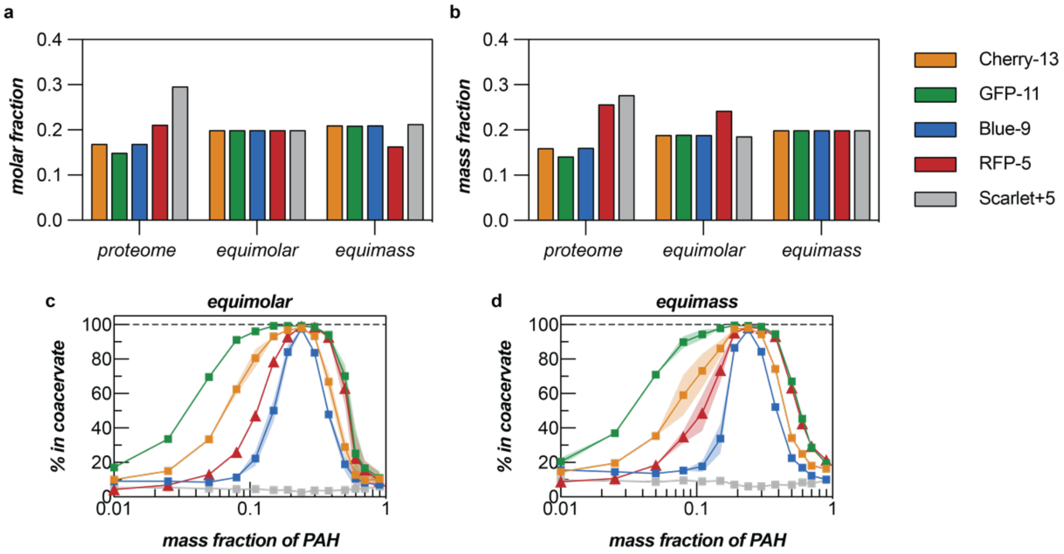


**Supplementary Figure 7**. (a) Molar and (b) mass fraction of proteins in the proteome-mimicking mixture, equimolar mixture, and equimass mixture. Partitioning analysis of (c) equimolar mixture and (d) equimass mixture.

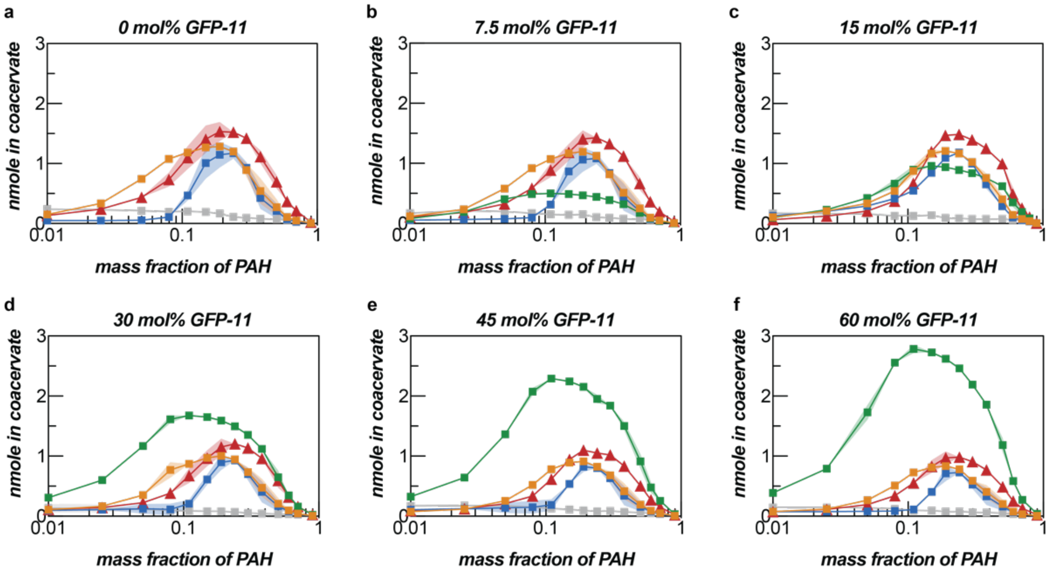


**Supplementary Figure 8**. Absolute partitioning analysis of proteome mimicking mixture with GFP-11 molar fraction of (a) 0%, (b) 7.5%, (c) 15%, (d) 30%, (e) 45%, and (f) 60%. The macromolecule concentration was fixed at 2.5 mg/mL but the percent of GFP was increased from 0-60% of the protein mixture.

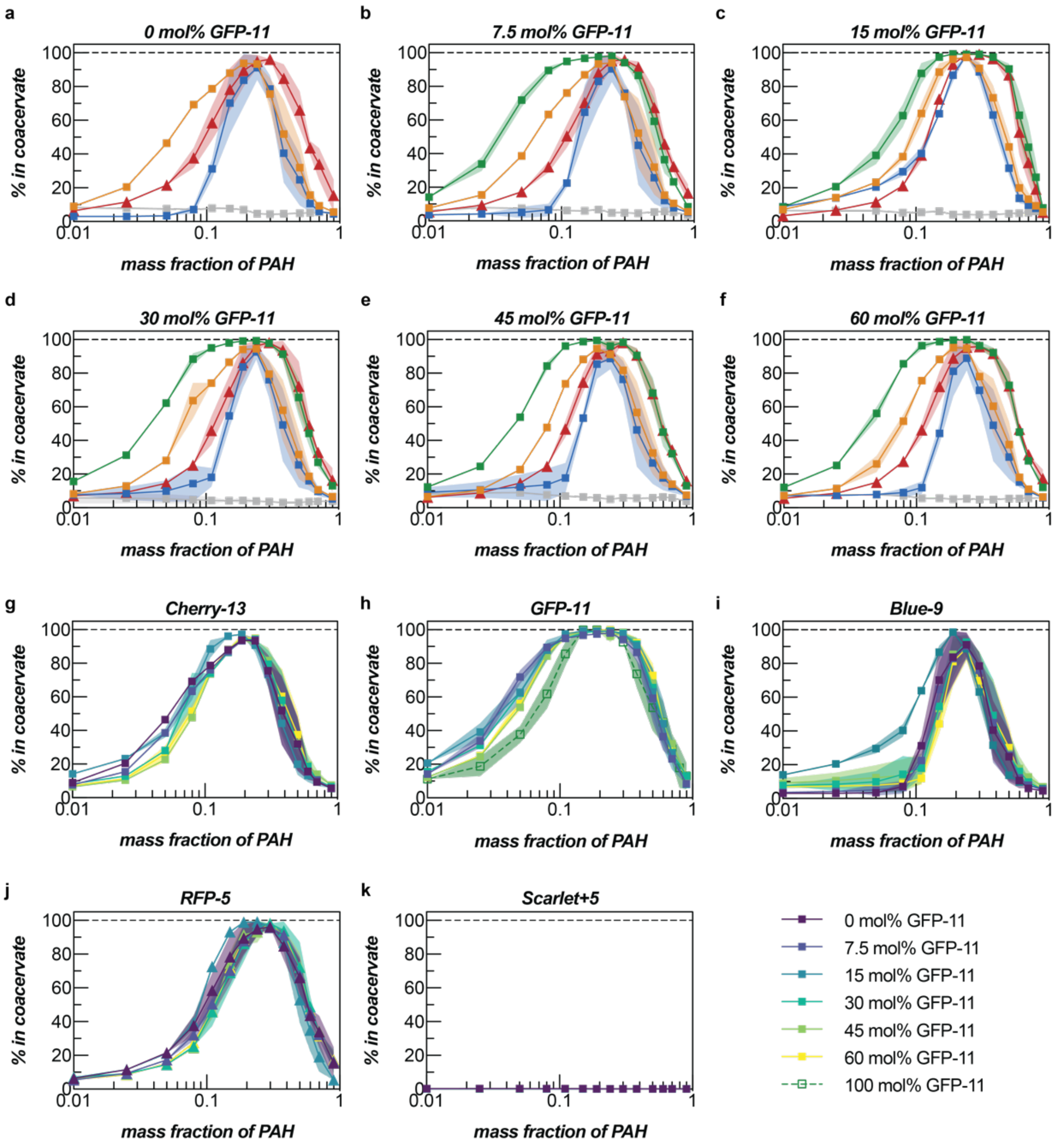


**Supplementary Figure 9**. Relative partitioning analysis of proteome mimicking mixture with GFP-11 molar fraction of (a) 0%, (b) 7.5%, (c) 15%, (d) 30%, (e) 45%, and (f) 60%. Relative partitioning analysis of (g) Cherry-13, (h) GFP-11, (i) Blue-9, (j) RFP-5, and (k) Scarlet+5 for all mass fraction conditions.

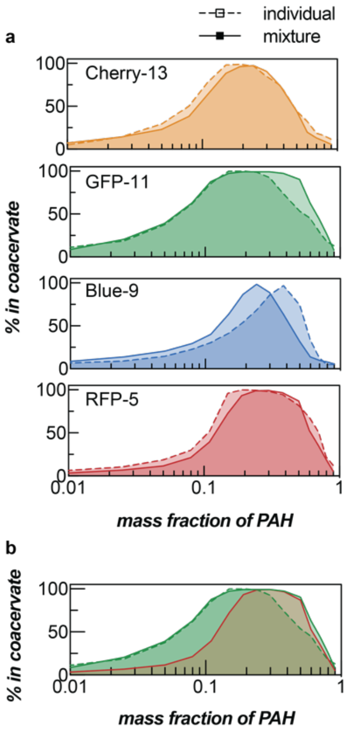


**Supplementary Figure 10**. (a) Partitioning comparison of individual proteins (dashed curves) and protein mixtures (solid lines). (b) Partitioning comparison among individual GFP-11 (green dashed line), GFP-11 in the mixture (green solid line) and RFP-5 in the mixture (red solid line). Partitioning curve of GFP-11 significantly broadened in the mixture, which overlaps with that of RFP-5 at higher PAH mass fractions.

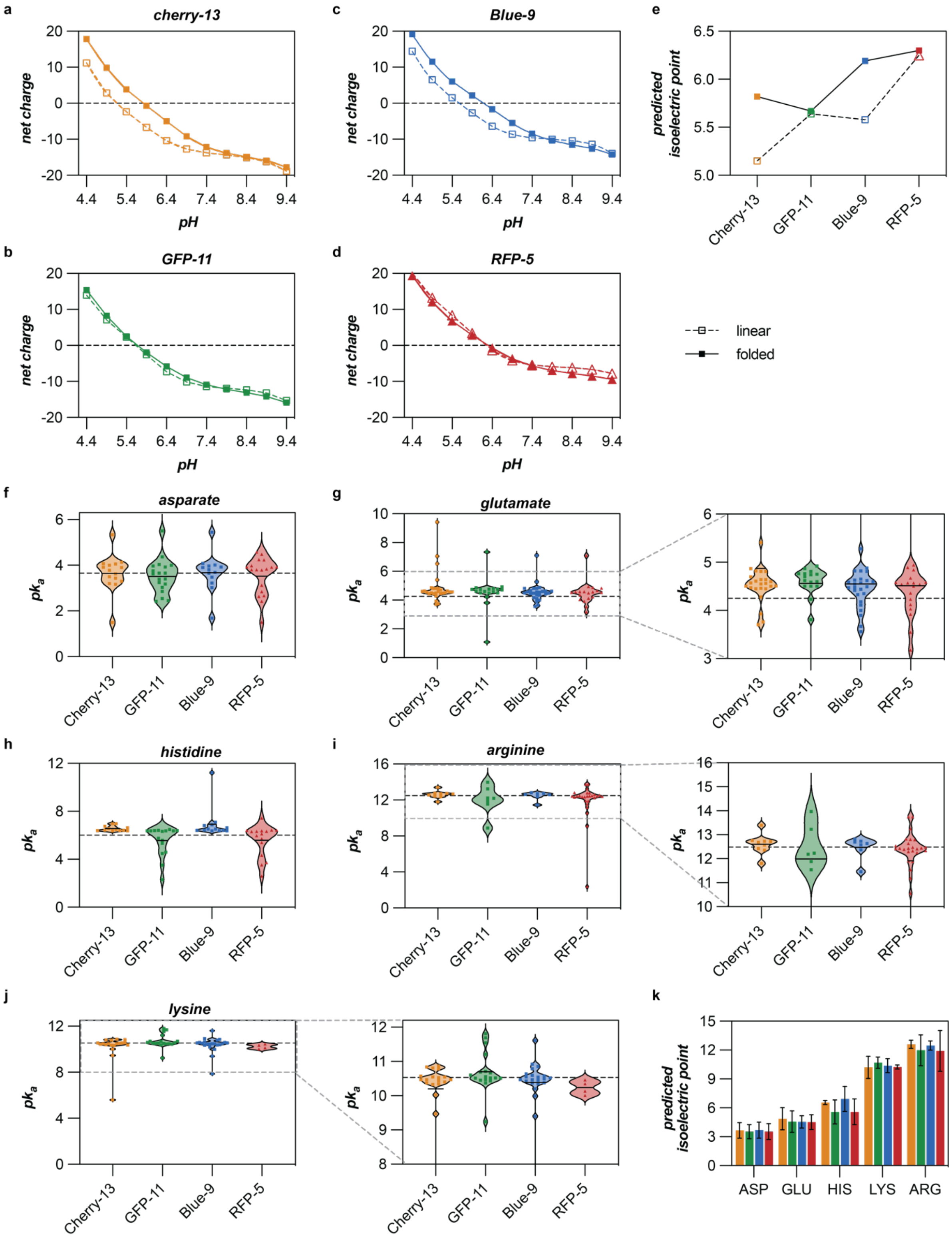


**Supplementary Figure 11**. Protein net charge and $pK_a$ values of residues predicted from linear sequence and PROPKA3. Net charge of (a) Cherry-13, (b) GFP-11, (c) Blue-9, and (d) RFP-5 based on theoretical $pK_a$ values of amino acids and linear sequence alone and based on 3D structure of the proteins determined by PROPKA3[18]. (e) Isoelectric point calculated based on the residue $pK_a$ prediction by PROPKA3. $pK_a$ values of all (f) aspartate, (g) glutamate, (h) histidine, (i) arginine, and (j) lysine residues on individual proteins predicted by PROPKA3. (k) Mean and standard deviation of $pK_a$ values of charged residues on individual proteins.

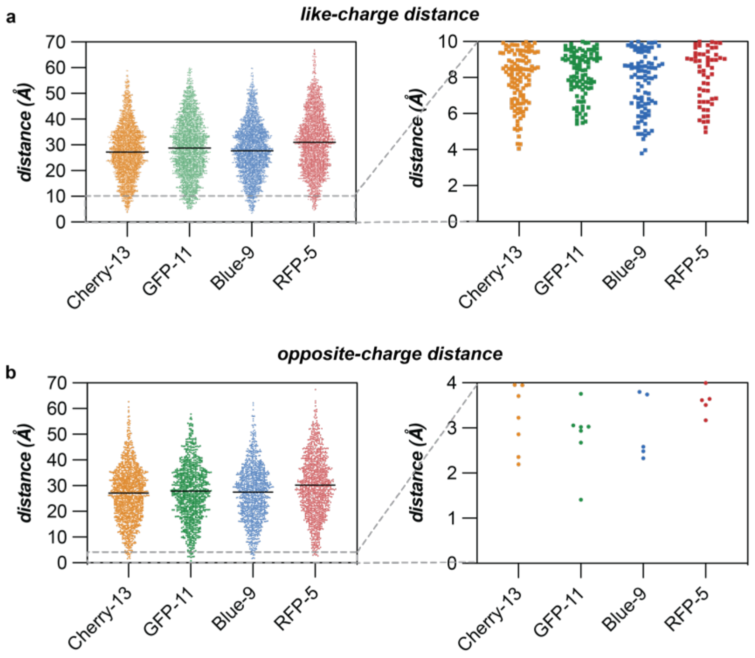


**Supplementary Figure 12**. Distance analysis of negatively charged residues. (a) Distances between oxygen atoms on carboxylic groups of all aspartate and glutamate residues. The lines in the plot indicate the mean distance, which correspond to 27.7 Å, 29.3 Å, 28.1 Å, and 31.4 Å for Cherry-13, GFP-11, Blue-9, and RFP-5 respectively. (b) Distances between negatively charged residues, aspartate and glutamate, and positively charged residues, arginine and lysine. Deprotonated oxygen atoms of aspartates and glutamates and protonated nitrogen atoms of arginines and lysines were selected to determine the distance between the charged residues. The lines in the plot indicate the mean distance, which corresponds to 27.2 Å, 27.8 Å, 27.5 Å, and 30.2 Å for Cherry-13, GFP-11, Blue-9, and RFP-5 respectively. In contrast to the distances between like-charges, more pairs of opposite-charges on GFP-11 were in close proximity, especially under 4 Å, the threshold distance for the favorable formation of salt bridges[23].

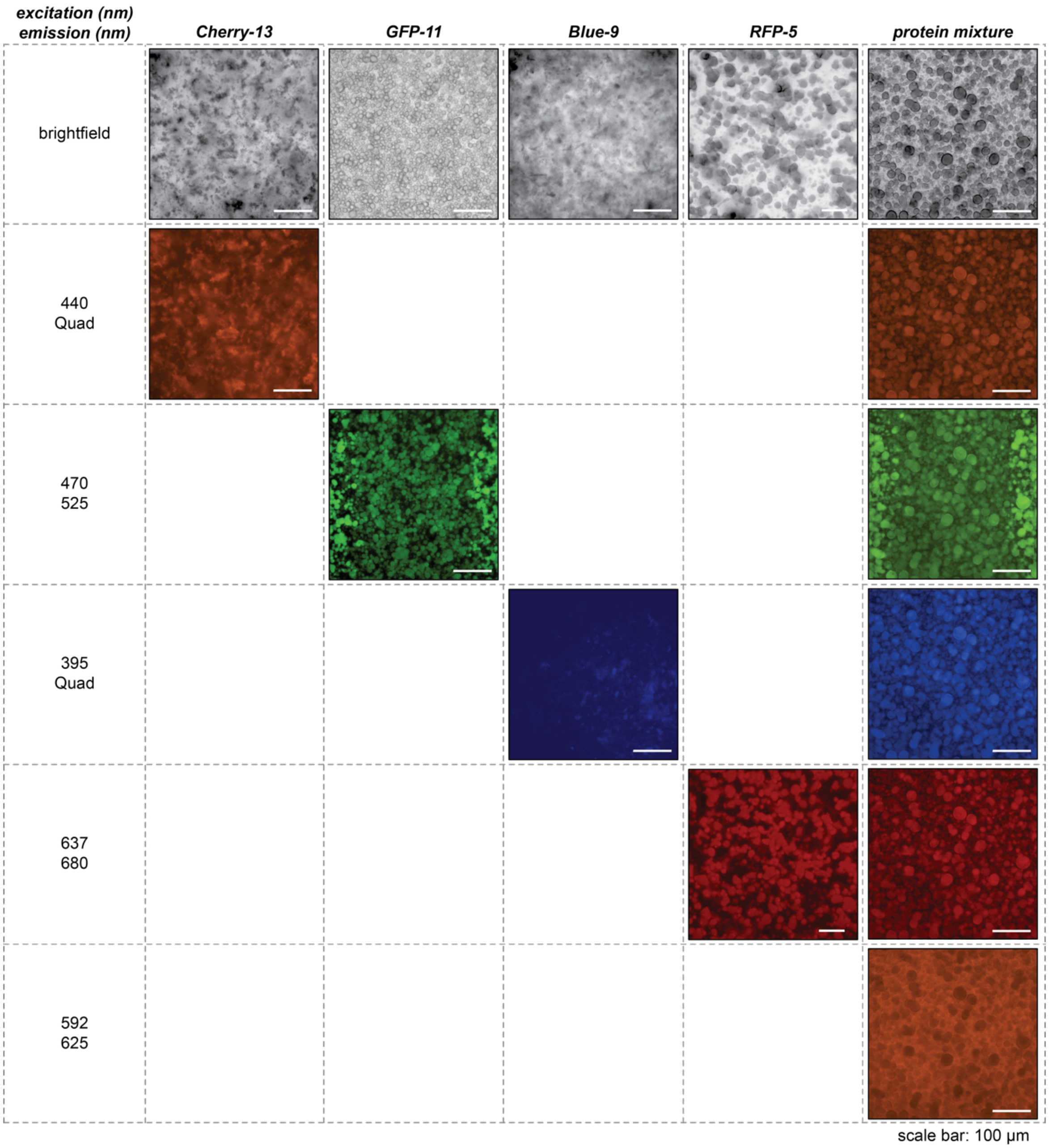


**Supplementary Figure 13**. Microscopic images of the coacervates of individual proteins and protein mixture. Quad emission filter passes wavelengths between 410 – 450 nm, 500 – 540 nm, 575 – 615 nm, and 665 – 770 nm. Complex coacervation was induced at total macromolecule concentration of 2.5 mg/mL, pH 7.4 with 0.19 mass fraction of PAH (0.475 mg/mL PAH + 2.025 mg/mL proteins). Noticeable differences in phase properties of coacervates between proteins were observed, with Cherry-13 and Blueberry-9 forming solid-like aggregates.

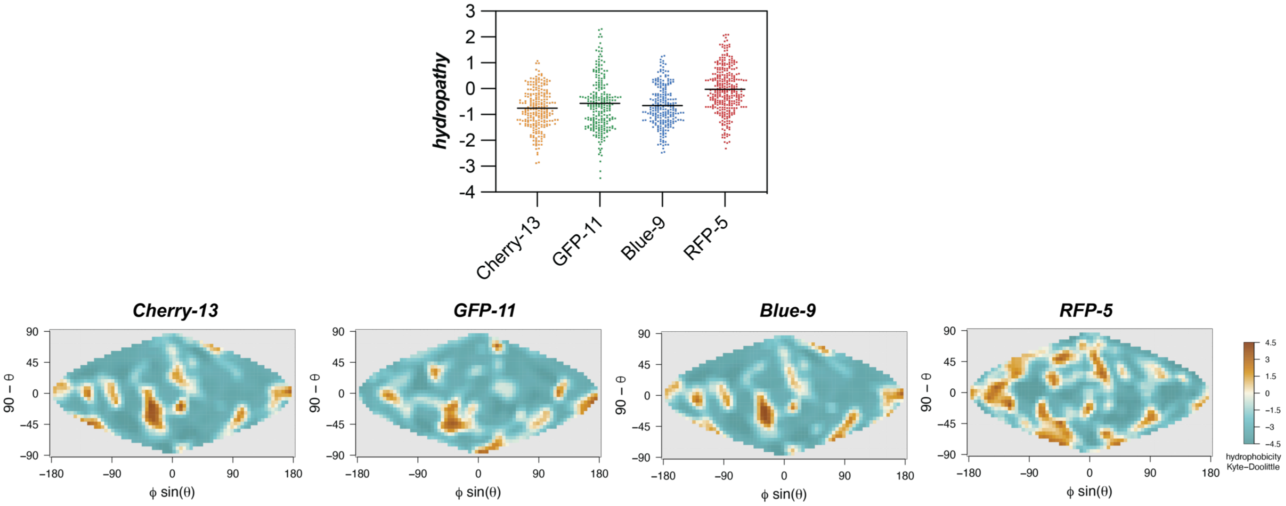


**Supplementary Figure 14**. Hydropathy values of residues in individual anionic proteins and 2D mapping of protein surface hydropathy. Hydropathy values were extracted from Kyte-Dolittle hydropathy plot[24]. 2D surface maps of the 3D structure of individual proteins were generated by SURFMAP[25]. The lines in the hydropathy plot indicate the mean hydropathy values of all residues on individual proteins which correspond to -0.75, -0.57, -0.65 and -0.02 for Cherry-13, GFP-11, Blue-9, and RFP-5 respectively.

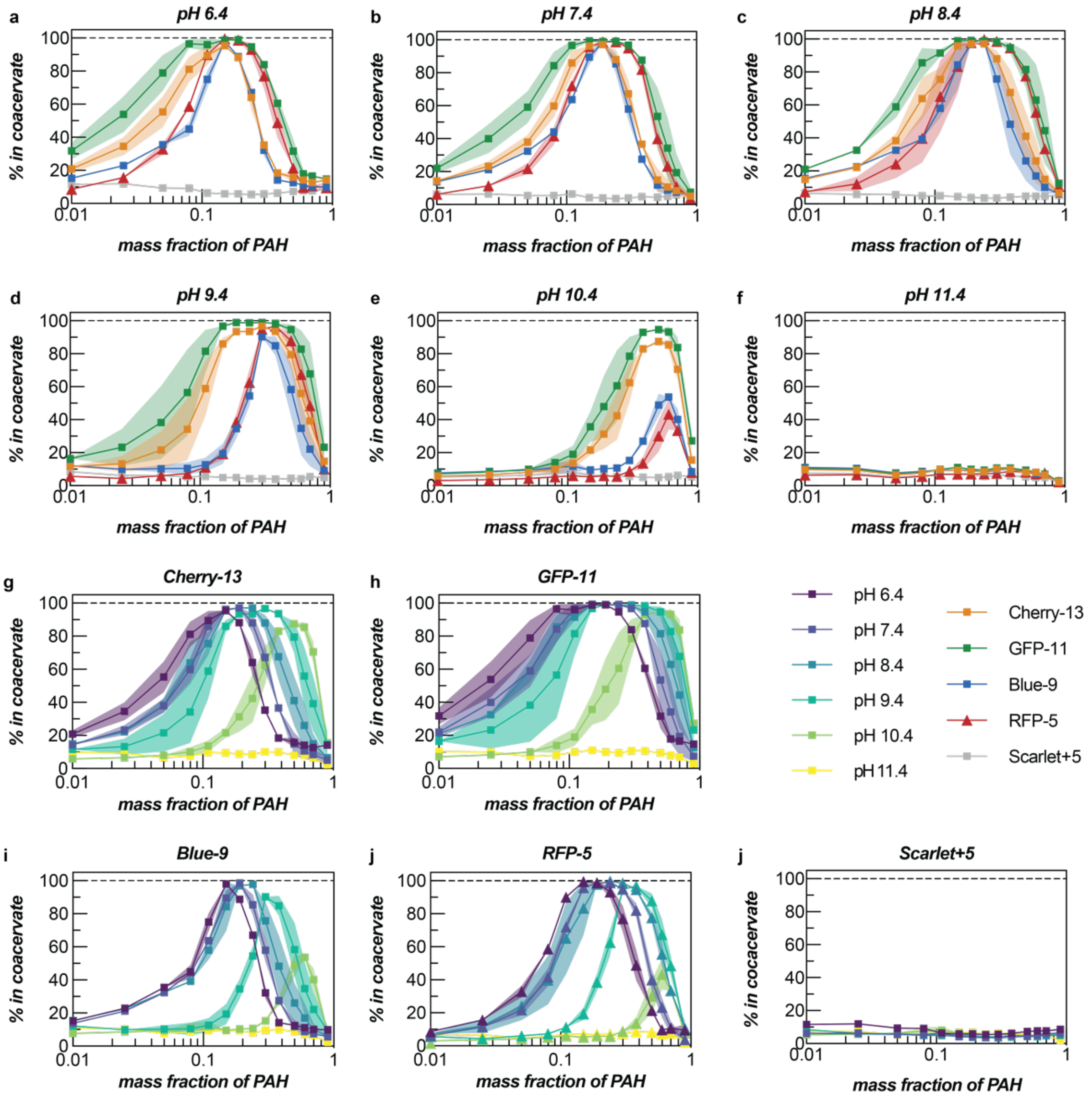


**Supplementary Figure 15**. Partitioning analysis of protein mixture at (a) pH 6.4, (b) pH 7.4, (c) pH 8.4, (d) pH 9.4, (e) pH 10.4, and (f) pH 11.4. Partitioning curves of all pH conditions of (g) Cherry-13, (h) GFP-11, (i) Blue-9, (j) RFP-5, and (j) Scarlet+5. Complex coacervation was induced at total macromolecule concentration of 2.5 mg/mL in 10 mM tris buffer adjusted to the specific pH.

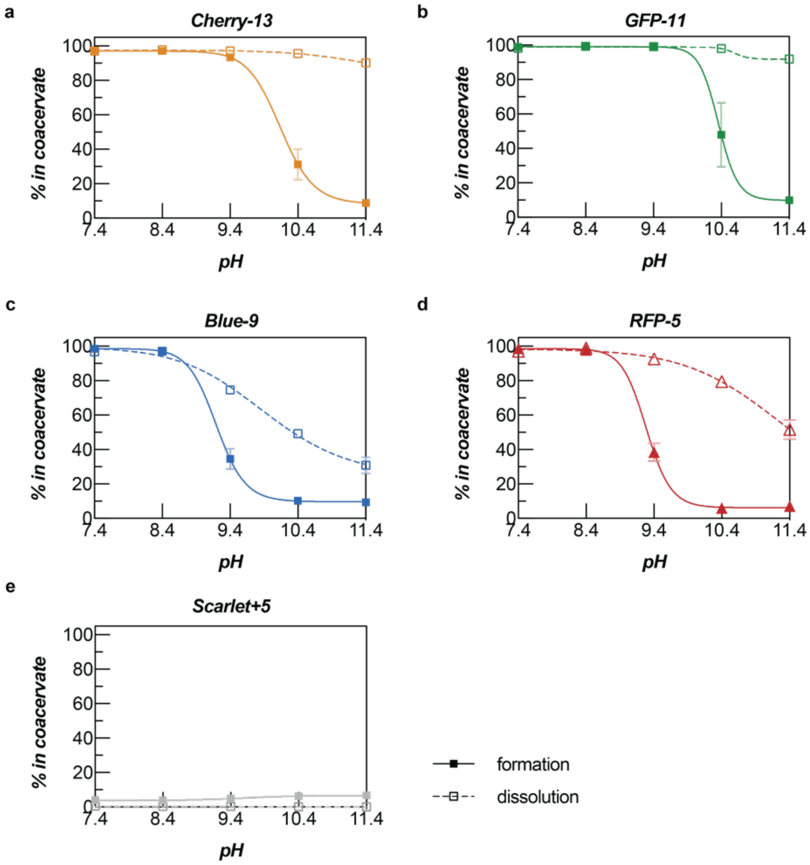


**Supplementary Figure 16.** Partitioning analysis of (a) Cherry-13, (b) GFP-11, (c) Blue-9, (d) RFP-5, and (e) Scarlet+5 in the protein mixture as a function of pH when the pH was varied either during coacervate formation or dissolution. Complex coacervation was induced with 2.025 mg/mL of the protein mixture and 0.475 mg/mL of a PAH solution (PAH mass fraction 0.19). For initial pH variation, 10 mM Tris-HCl buffer, protein solutions, and polymer solution were titrated to the desired pH separately prior to mixing. For pH dependent dissolution of the coacervates, the dense phase isolated by centrifugation was resuspended with 100 µL of 10 mM Tris-HCl buffer with pH varying from 6.4 to 11.4 by gentle pipetting and bath sonication (5 s on, 5 s off, repeated 3 times). The partially resuspended dense phase was incubated for 30 min at room temperature, followed by separation and analysis.

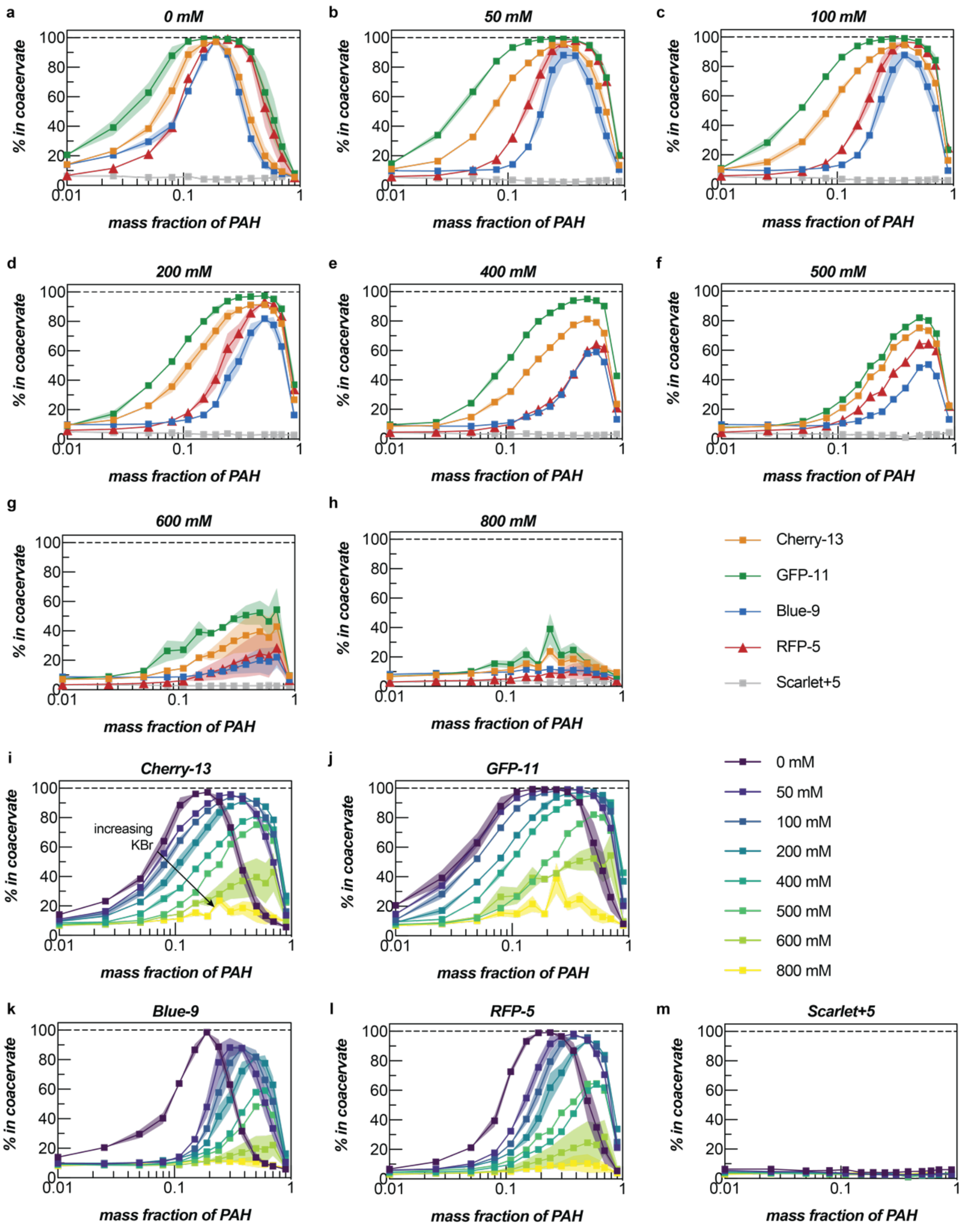


**Supplementary Figure 17.** Partitioning analysis of protein mixture at (a) 0 mM, (b) 50 mM, (c) 100 mM, (d) 200 mM, (e) 400 mM, (f) 500 mM, (g) 600 mM, and (h) 800 mM KBr. Partitioning curves of all ionic strength conditions of (i) Cherry-13, (j) GFP-11, (k) Blue-9, (l) RFP-5, and (m) Scarlet+5. Complex coacervation was induced at total macromolecule concentration of 2.5 mg/mL at pH 7.4.

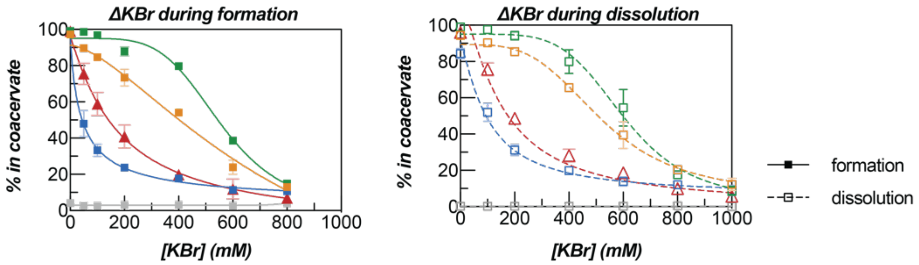


**Supplementary Figure 18.** Partitioning analysis of proteins in the protein mixture during (a) formation and (b) dissolution of the coacervates at various ionic strengths. Complex coacervation was induced at 2.025 mg/mL of the protein mixture and 0.475 mg/mL of a PAH solution (PAH mass fraction 0.19). For initial ionic strength variation, the KBr concentration in 10 mM Tris buffer, protein solution, and polymer solution was adjusted prior to mixing. For ionic strength dependent dissolution of the coacervates, the dense phase isolated by centrifugation was resuspended with 100 µL of a KBr solution, with KBr concentration ranging from 100 mM to 1000 mM, by gentle pipetting and bath sonication (5 s on, 5 s off, repeated 3 times). The partially resuspended dense phase was incubated for 30 min at room temperature, followed by separation and analysis.

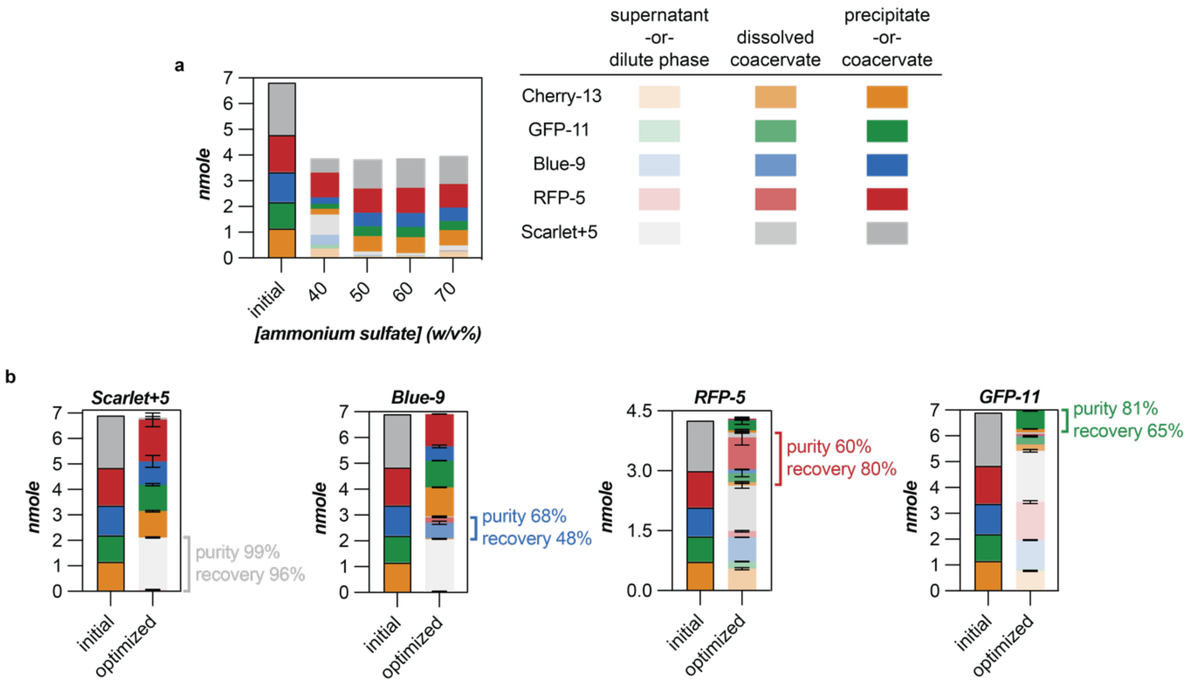


**Supplementary Figure 19**. Quantity of individual proteins in the initial mixture and the absolute recovery (a) for ammonium sulfate precipitation and (b) for complex coacervation with optimized conditions. The bars are colored based on the protein and shaded based on where the protein was recovered from (*e.g.* lighter bars correspond to the dilute phase or supernatant). No obvious selectivity was observed between the proteins with ammonium sulfate precipitation and the maximum recovery of the proteins from both phases was around 57%, which is significantly lower than that of complex coacervation. The sum of proteins in each phase was nearly 100% at optimized purification conditions by complex coacervation.
